# Structural and Biochemical Rationale for Enhanced Spike Protein Fitness in Delta and Kappa SARS-CoV-2 Variants

**DOI:** 10.1101/2021.09.02.458774

**Authors:** James W. Saville, Dhiraj Mannar, Xing Zhu, Shanti S. Srivastava, Alison M. Berezuk, Jean-Philippe Demers, Steven Zhou, Katharine S. Tuttle, Inna Sekirov, Andrew Kim, Wei Li, Dimiter S. Dimitrov, Sriram Subramaniam

## Abstract

The Delta and Kappa variants of SARS-CoV-2 co-emerged in India in late 2020, with the Delta variant underlying the resurgence of COVID-19, even in countries with high vaccination rates. In this study, we assess structural and biochemical aspects of viral fitness for these two variants using cryo-electron microscopy (cryo-EM), ACE2-binding and antibody neutralization analyses. Both variants demonstrate escape of antibodies targeting the N-terminal domain, an important immune hotspot for neutralizing epitopes. Compared to wild-type and Kappa lineages, Delta variant spike proteins show modest increase in ACE2 affinity, likely due to enhanced electrostatic complementarity at the RBD-ACE2 interface, which we characterize by cryo-EM. Unexpectedly, Kappa variant spike trimers form a novel head-to-head dimer-of-trimers assembly, which we demonstrate is a result of the E484Q mutation. The combination of increased antibody escape and enhanced ACE2 binding provides an explanation, in part, for the rapid global dominance of the Delta variant.

## Introduction

In March 2021, genomic sequencing of SARS-CoV-2 samples in Maharashtra, India revealed increased prevalence of E484Q, L452R, and P681R co-mutation in the Spike glycoprotein (S protein)^1, 2, 3^. This variant was called the “double mutant” by the global news media and was later designated as lineage B.1.617.1 and then the Kappa variant of interest by the World Health Organization^4^. The Kappa (B.1.617.1) lineage is a sub-lineage of the B.1.617 lineage, which is defined by L452R and P681R co-mutation. By early April 2021, the Kappa variant accounted for approximately 35% of all sequenced cases in India, which coincided with the start of a rise in daily COVID-19 cases (Figure 1A,B) ^1, 2, 3^. In addition to the B.1.617 and B.1.617.1 variants, variants B.1.617.2 and B.1.617.3 saw increased prevalence in March 2021. While the B.1.617 and B.1.617.3 variants never exceeded 5% of sequenced cases in India, the B.1.617.2 variant (now designated as the Delta variant of concern) rapidly became dominant in India within months. Given the precursory identification of the Kappa variant, it was initially thought to be the major contributor to the increased COVID-19 case numbers in India. However, retrospective analyses show the rapid dominance of the Delta variant to better coincide with the exponential surge of COVID-19 cases in India (Figure 1A,B).

**Figure 1:**
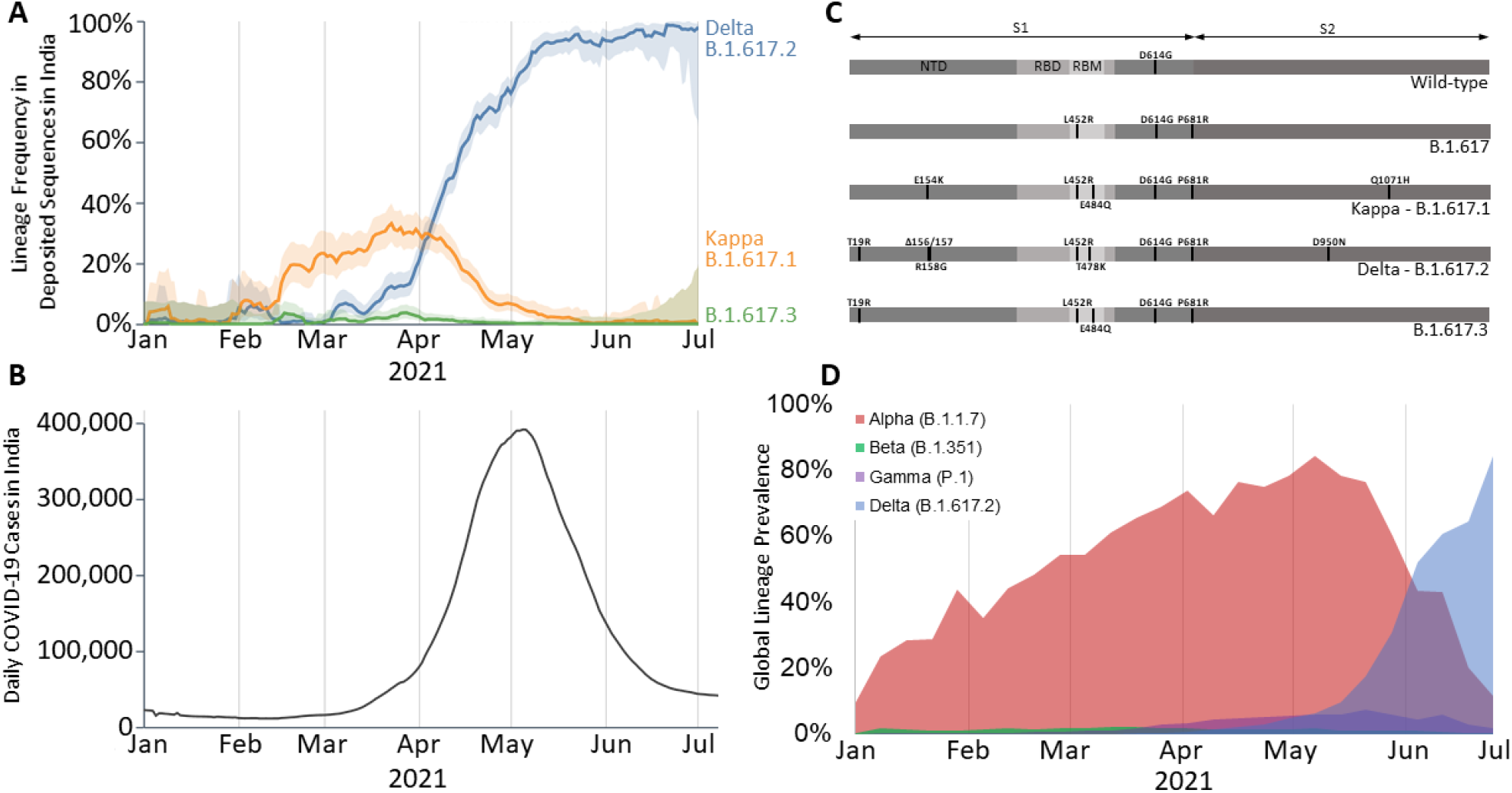
Emergence and prevalence of the Delta and Kappa SARS-CoV-2 lineages. **(A)** Delta (B.1.617.2), Kappa (B.1.617.1), and B.1.617.3 lineage frequency in India from January to July 2021. Deposited sequences were downloaded from the GISAID initiative. The 95% confidence interval is plotted as the lighter shaded border around each line. This figure panel was generated using the outbreak.info project^3^. **(B)** Daily confirmed COVID19 cases in India as curated by the outbreak.info project^3^. **(C)** SARS-CoV-2 amino acid sequence box plots for the wild-type (D614G) and B.1.617 sub-lineages. The abbreviations are NTD: N-terminal domain, RBD: Receptor-binding domain, and RBM: Receptor binding motif. **(D)** Global prevalence for the Alpha, Beta, Gamma, and Delta lineages of SARS-CoV-2 from January to July 2021. The sequence data were downloaded from the GISAID initiative and plotted as the percentage of the global prevalence for each week^1,2^.

Following the emergence of the Delta variant in India, it rapidly spread globally and in early June 2021, it overtook the Alpha (B.1.1.7) variant to become the major SARS-CoV-2 lineage worldwide (Figure 1D)^1, 2, 3^. The Kappa variant, despite harbouring multiple mutations in the S protein that have been demonstrated to increase viral fitness, has not significantly spread globally, with only a few travel-linked clusters reported^5, 6, 7^. The exact mutations present within the Kappa and Delta S proteins have been reported variously, suggesting a degree of variability in their classification^8, 9, 10, 11, 12^. Within the receptor-binding domain (RBD) of the S protein, both the Kappa and Delta variants share an identical substitutional mutation (L452R) with the previously emerged variants of interest B.1.427/429 (Epsilon)^1,2,4,8^. The S protein E484Q mutation in the Kappa variant was similarly found in the Beta (B.1.351) and Gamma (P.1) lineages, where residue 484 is mutated to lysine (E484K) ^1,2^. The Delta variant contains a novel T478K mutation within the RBD - that is not present in previous variants of concern - with uncharacterized effect. After the RBD, the NTD is the second highest-targeted S protein domain by SARS-CoV-2 convalescent sera antibodies^13^. The NTD is comprised of five loops (called N1-5; N1: residues 14-26, N2: residues 67-79, N3: residues 141-156, N4: residues 177-189, and N5: residues 246-260), with N1, N3, and N5 loops dubbed an “NTD neutralization supersite” for the propensity of neutralizing antibodies to target this location^14^. While the Kappa variant contains only a single mutation (E154K) in N3, the Delta variant contains a mutation in N1 (T19R) and multiple mutations in N3 (Δ156/157, R158G). Finally, the P681R mutation (present in all B.1.617 lineages) immediately precedes the furin cleavage consensus sequence (682-RRSR/SVA-688), with initial reports suggesting that the P681R mutation enhances cleavage, and consequently S protein post-fusion transition^11,15,16^. How the combinatorial effect of the Delta and Kappa S protein mutations contribute towards increased viral fitness remains superficially characterized relative to previously emerged variants of concern (Alpha, Beta, Gamma).

Herein, we report cryo-electron microscopy (cryo-EM) structures of Delta and Kappa variant S protein trimers - both in the unbound state and in complex with the ACE2 receptor - to gain insight into how their mutations underlie changes in ACE2 binding and antibody neutralization escape. Through our experiments, we uncovered an unprecedented dimerization phenomenon for the Kappa variant S protein with - as yet -unknown biological implications. We go onto report two additional structures of S proteins containing novel substitutions at position 484 (I484 and A484) to dissect the chemical nature of this dimerization event.

## Results

### Antibody Evasion by Delta and Kappa Spike proteins

To assess the antibody evasive properties of the Kappa and Delta spike protein mutations, we performed antibody binding and neutralization studies using purified variant spike ectodomains (Figure S1) and variant pseudo-typed viruses respectively. First, a panel of six previously characterized monoclonal antibodies targeting epitopes in the spike RBD or NTD were assessed (Figure 2A)^17, 18, 19, 20, 21, 22^. Interestingly, we observed enhanced potency of ab1 for both Kappa and Delta variant spikes relative to wild-type, despite the presence of the Kappa T478K mutation which falls within the ab1 footprint. Analysis of the RBD-ab1 interface reveals close proximity of the ab1 amino-terminal glutamic acid to position 478 within the RBD (Figure S2). Thus, the enhanced potency of ab1 for the Delta variant is likely explained by the formation of an additional electrostatic interaction and/or salt bridge. We have previously demonstrated the sensitivity of ab8 and S2M11 to the E484K mutation, wherein both antibodies were fully escaped^23^. While total escape of ab8 was achieved by the E484Q-bearing Kappa variant spike, binding and neutralization of S2M11 was attenuated but not abolished (Figure 2A), demonstrating the relative sensitivities of these antibodies to mutation of E484. Although the S309 footprint does not include any Delta or Kappa RBD mutational positions, we observed reduced binding and neutralizing potencies for both of these variant spikes relative to wild-type, which corroborates recent results^23^. The Kappa variant spike harbours mutations that fall within the footprints of both NTD-directed antibodies tested (4A8 and 4-8), and accordingly demonstrated total escape (Figure 2B). The Delta spike contains NTD mutations within the footprint of 4-8 but not 4A8 and similarly evaded both antibodies (Figure 2B), suggesting both direct and indirect mechanisms of evasion. Second, we evaluated the impact of Kappa and Delta spike protein mutations on neutralization by polyclonal preparations of human antibodies. To this end, we utilized sera from a set of patients with varying vaccination statuses and COVID-19 histories (Figure S13) in pseudo-typed virus entry assays. We observed different degrees of attenuated neutralization potencies for both variants when compared to wild-type and observed a statistically significant decrease in potency for the Delta variant (Figure 2C, p=0.02).

**Figure 2:**
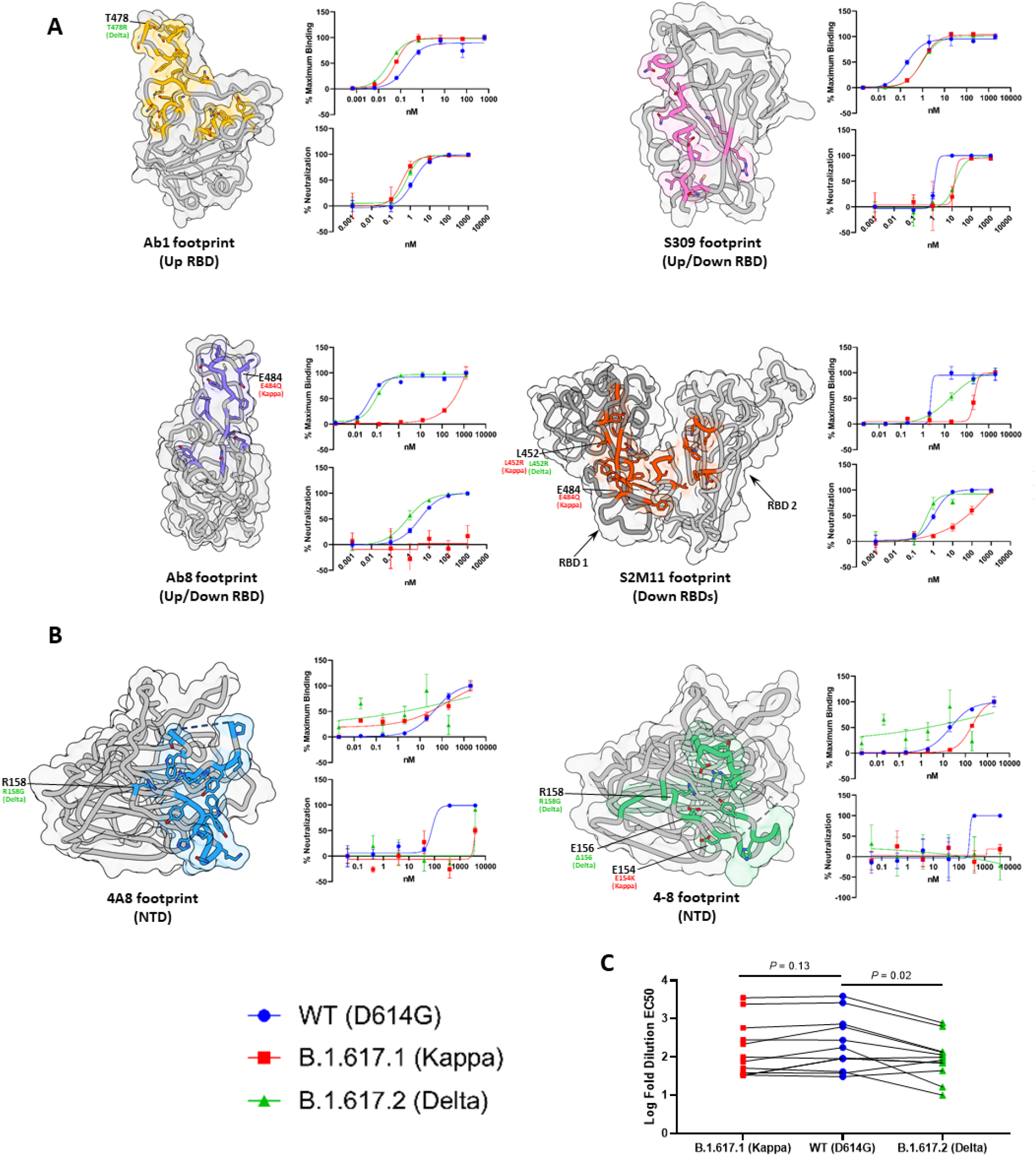
Antibody evasion by Delta and Kappa variants. **(A)** Antibody-binding footprints, antibody binding as assessed by ELISA, and pseudovirus neutralization for four anti-RBD antibodies (Ab1, Ab8, S309, and S2M11). **(B)** As in (A) but for two anti-NTD antibodies (4A8 and 4-8). **(C)** Log fold dilutions of sera conferring 50% neutralization (EC_50_) of pseudovirus harbouring wild-type, Kappa, or Delta spikes, from patients post COVID19 and/or vaccination (n = 12). Patient-derived sera sample information and raw pseudovirus neutralization data are presented in Figure S13.

### Delta Variant S Protein Mutations Moderately Increase ACE2 Affinity as a Consequence of Enhanced Electrostatic Complementarity at the RBD-ACE2 Interface

As demonstrated in previously emerged SARS-CoV-2 variants of concern (Beta, Gamma), evolution of the S protein seems to select for combinations of mutations that balance both antibody evasion and ACE2 binding affinity ^23, 24, 25^. Having observed monoclonal and polyclonal antibody evasion for the Delta and Kappa variants (Figure 2A,B), we next sought to assess the impact of these mutations on ACE2 affinity. Biolayer interferometry (BLI) analysis of trimeric variant S proteins revealed limited (no change) and moderate (∼2-fold) enhancements in ACE2 affinity for the Kappa and Delta variants respectively (Figure 3C, Figure S3). To investigate a structural rationale for the observed changes - or lack thereof - in ACE2 binding affinity, we solved the cryo-EM structures of the Kappa and Delta variant spike trimers in complex with ACE2 (Figure 3A, Figure S4, Figure S5).

**Figure 3:**
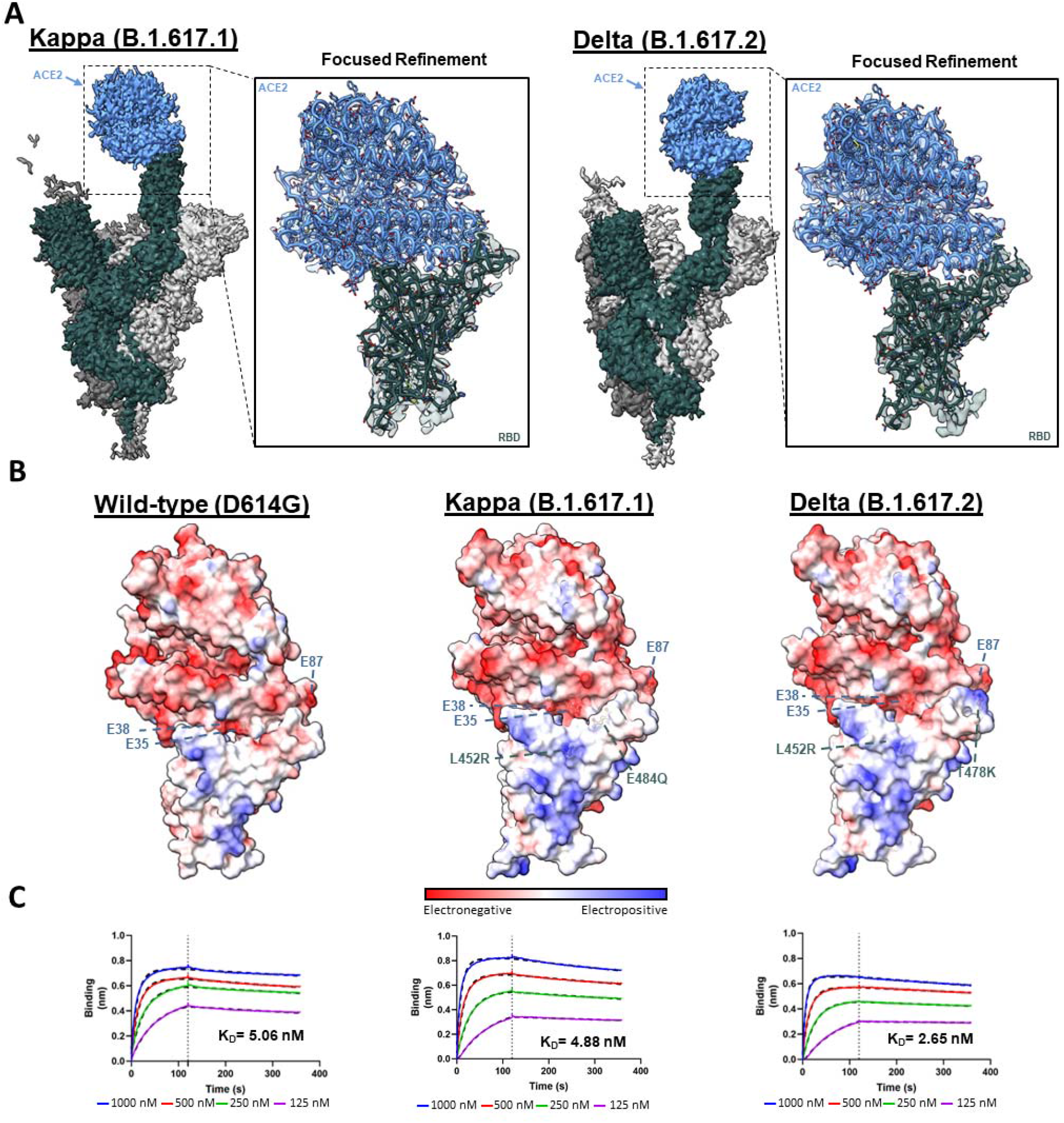
Structural and biophysical effects of Kappa and Delta S protein mutations on ACE2 binding. **(A)** The global and focus-refined cryo-EM structures for the Kappa and Delta variant S proteins in complex with ACE2. **(B)** Electrostatic surface potentials for the wild-type, Kappa, and Delta S protein– ACE2 complexes. We previously reported the structure of the wild-type S protein–ACE2 complex and we use this model here to generate the wild-type electrostatic surface^23^. **(C)** Biolayer interferometry (BLI) sensorgrams for the wild-type, Kappa, and Delta variant S protein–ACE2 binding interaction. The S protein concentrations utilized are: 1000 nM (blue), 500 nM (red), 250 nM (green), 125 nM (magenta). The *k*_*on*_ and k_off_ rate constants are presented in Figure S3.

The focus-refined atomic structure of the Kappa variant S protein in complex with ACE2 reveals limited structural changes at the RBD-ACE2 interface (Figure 3A). The Kappa variant E484Q mutation results in the loss of an electrostatic interaction between residue E484 and residue K31 within ACE2, likely resulting in a weaker interaction at this site. However, the enhanced electrostatic complementarity afforded by the accompanying L452R mutation, as described previously, may present a compensatory mutation accounting for the lost E484–K31 interaction (Figure 3B)^23^. The combination of these two opposing mutations, one diminishing ACE2 affinity (E484Q), and the other increasing ACE2 binding (L452R), is consistent with the unchanged overall affinity of the Kappa S protein–ACE2 binding interaction. Precedence for compensatory mutations towards increasing ACE2 affinity while decreasing antibody binding has been reported for the N501Y and K417N/T mutational combinations found in the Beta and Gamma variants^23, 24, 25, 26^. Interestingly, the overall unchanged ACE2 affinity of the Kappa variant S protein stands in contrast to the majority of previously characterized SARS-CoV-2 variants of concern (Alpha, Beta, Gamma)^27, 28, 29, 30, 31^.

As for the Kappa variant, the Delta variant S protein–ACE2 complex focus-refined cryo-EM structure reveals limited sidechain rearrangement at the RBD-ACE2 interface (Figure 3A). The Delta variant lacks the E484Q substitution which preserves the E484–K31 electrostatic interaction, while the common L452R mutation may increase ACE2 binding by enhancing electrostatic complementarity^23^. Further, the Delta variant lysine substitution at position 478 (T478K) extends its positively charged sidechain towards an electronegative region on ACE2 (centered at position E87) (Figure 3B). Therefore, the combination of enhanced electrostatic complementarity afforded by the L452R and T478K Delta variant substitutions likely accounts for the moderate increase in ACE2 affinity.

### Cryo-EM Structure of the Kappa Variant S Protein Reveals a Head-to-Head Dimerization Phenomenon

We next aimed to evaluate the effect of Kappa and Delta S protein mutations on spike conformation and quaternary structure. We solved cryo-EM structures of the Kappa and Delta spikes at global average resolutions of 3.16 and 2.25 Å respectively (Figures S6, S7, Table S1). While the cryo-EM 3D reconstruction of the Delta spike reveals no large-scale structural changes (Figure S11), the global reconstruction of the Kappa variant S protein uncovers a previously unreported head-to-head dimer-of-trimers phenotype (Figure 4A). This dimer is mediated by RBD-RBD contacts between two S protein trimers, with one trimer offset at a 12° angle. This angle is a result of slightly asymmetric binding at each RBD-RBD contact. Focused refinement of the dimer-of-trimers interaction interface reveals an extensive binding interface involving all six RBDs (Figure 4B). Each RBD interacts with two RBDs in the opposite trimer via two distinct interfaces, herein refer to as RBD1 and RBD2 (Figure 4C,D). Interactions stabilizing the RBD1 interface are primarily mediated by van der Waals interactions and hydrophobic contacts between residues across this interface. Additionally, homo-asparagine-asparagine and glutamine-glutamine hydrogen bonds at positions 440 and 506, respectively, and a backbone carbonyl oxygen– amide hydrogen bond between residues 372 and 445 further contribute to the RBD1 interface (Figure 4C).

**Figure 4:**
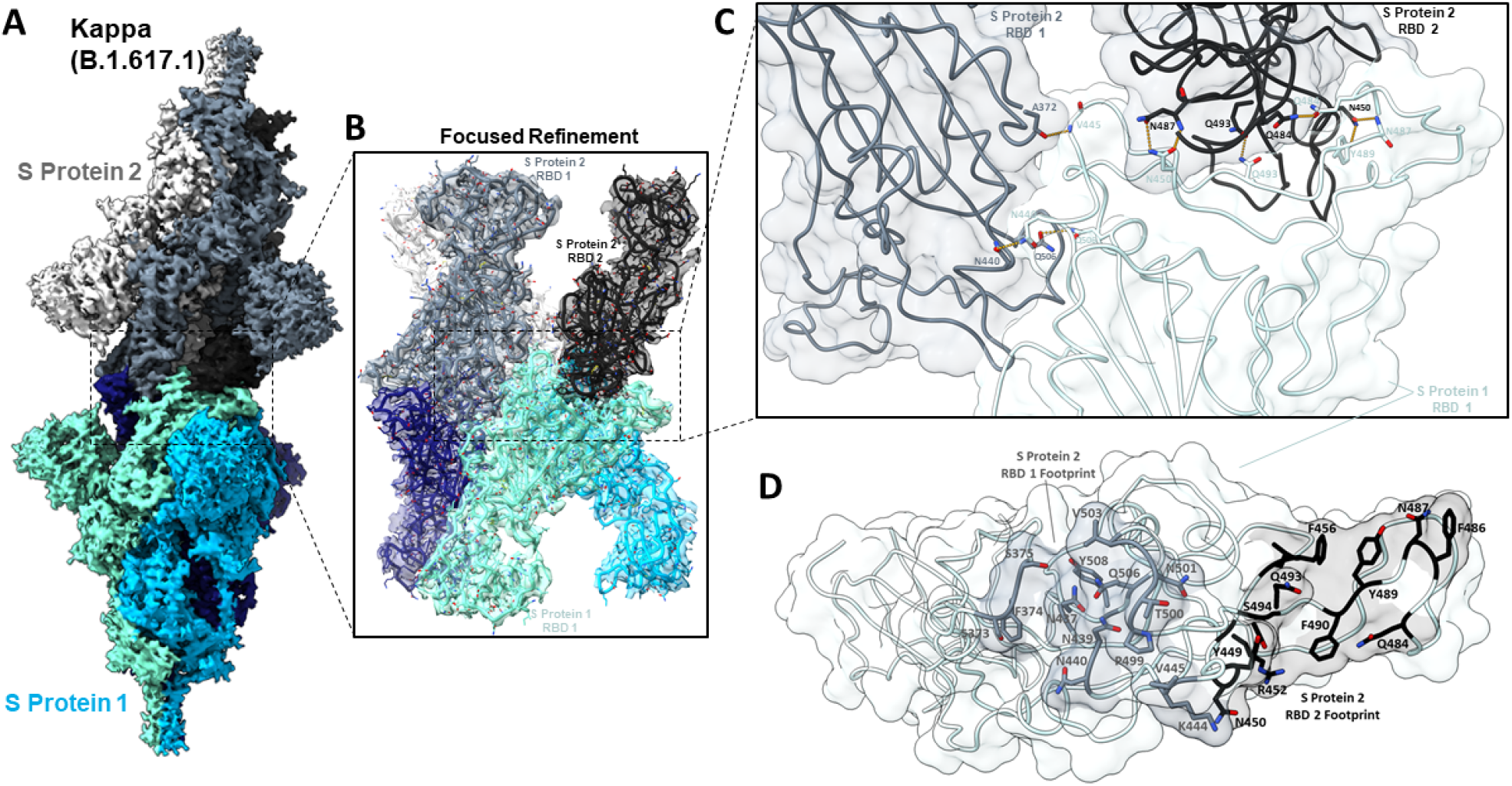
The Kappa variant S protein exhibits a novel dimer-of-trimers phenotype. **(A)** Side view of the global cryo-EM density map of the Kappa variant dimer-of-trimers complex. One trimer (S Protein 1, bottom) is displayed in various shades of blue, and the other trimer in grayscale (S Protein 2, top). **(B)** Focus-refined cryo-EM density map and fitted atomic model at the dimerization interface. **(C)** Detailed view of the molecular interface between two RBDs in the grayscale trimer (top) and a single RBD in the blue shaded trimer (bottom). For amino acids involved in inter-residue hydrogen bonding, the backbone and sidechain atoms are displayed. Hydrogen bonds are indicated by yellow dashed lines. **(D)** The top-down view of the region in panel (C) is shown. The binding footprints of RBD1 and RBD2 are indicated by complementary shading and displaying the sidechain atoms of footprint residues.

The second RBD-RBD interface (RBD2), comprises multiple doubly-hydrogen bonded residues, between residues N487-N450 and between residues N450-Y489/N487 (Figure 4C,D). Additionally, two homo-glutamine-glutamine hydrogen bonds are present between residues Q493-Q493 and Q484-Q484 from each trimer. This latter interaction at position 484 is of particular interest as it is uniquely mutated from glutamic acid to glutamine (E484Q) in the Kappa variant. Given this unique substitution and the unique dimer-of-trimers phenotype seen only for the Kappa variant spike (and not for Alpha, Beta, Gamma, Epsilon, and various RBD-mutated spike trimers), we identified position 484 as likely being crucial for S protein dimerization^23^.

### Reduced Charge Repulsion and Additional Sidechain Contacts at Position 484 Result in a Dimer-of-Trimers Phenotype

Having identified that residue identity at position 484 likely affects head-to-head S protein oligomerization, we aimed to further probe the chemical properties at 484 that mediate this dimerization. A focused view of the Q484-Q484 hydrogen bond (Figure 5A) shows the bond to be “sandwiched” by proximal bulky F490 aromatic sidechains. We therefore hypothesized that charge neutrality at position 484 (as seen in the Q484, but not E484 or the recently emerged K484 S proteins) may be sufficient to reduce charge-charge repulsion at this site and therefore allow dimerization. To test this, we performed site-directed mutagenesis to substitute an alanine at position 484 (Q484A) in the Kappa S protein, purified the trimer, and performed structural studies. The cryoEM reconstruction of Q484A spikes revealed no evidence of dimer-of-trimer assemblies, consistent with our previous results for wild-type and other variant of concern (VoC) spikes^23^ (Figure 5B). We next hypothesized that the homo-glutamine hydrogen bond conferred by the Q484 sidechain provided an additional contact critical for dimer formation. Accordingly, we introduced the amino acid isoleucine at position 484, which possesses a branched aliphatic sidechain capable of providing hydrophobic packing contacts. The cryo-EM reconstruction yielded a dimer-of-trimers phenotype for the Kappa + Q484I S protein variant, yet, with a reduced number of picked particles comprising the dimer class (46%), relative to the original Kappa variant with Q484 (74%) (Figure 5B,C). The oligomerization state of S proteins harbouring charged residues at 484 (E484, K484), along with the Q484A and Q484I mutations demonstrate that abrogation of charge at position 484 is necessary but not sufficient to permit dimerization. Rather, a combination of charge neutralization and additional contacts enabled by sidechains at position 484 is required for S protein dimerization.

**Figure 5:**
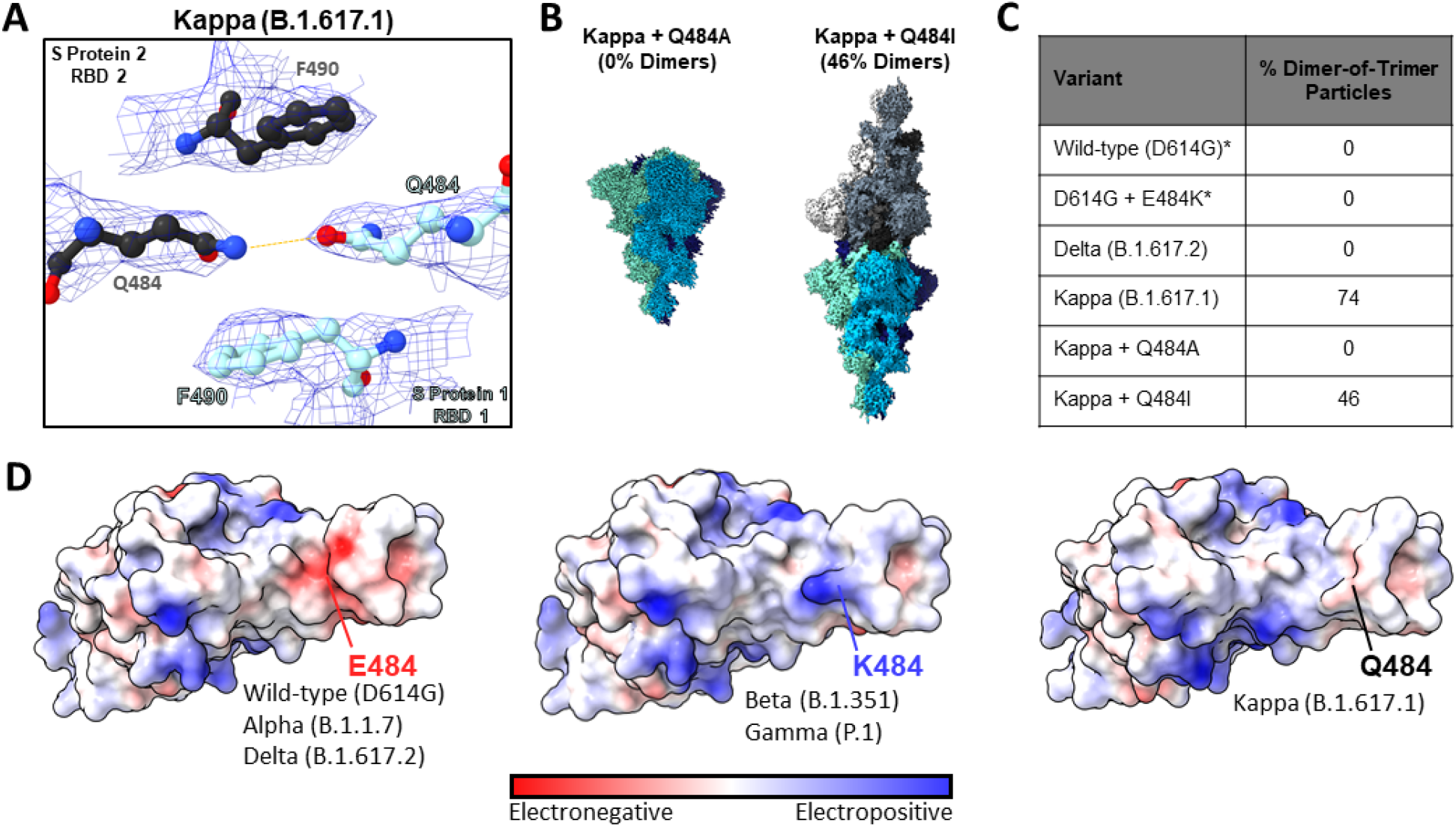
Impact of residue identity at position 484 on S protein oligomerization. **(A)** Detailed view of the Kappa (B.1.617.1) dimer-of-trimers cryo-EM density map and fitted model at the Q484-Q484 interaction site. The hydrogen bond formed between Q484 residues located in different S ectodomain trimers (S protein 2 RBD 2 in black, and S protein 1 RBD 1 in cyan) is indicated with a yellow dashed line. **(B)** Cryo-EM density maps for the Kappa + Q484A and Kappa + Q484I mutated S proteins. **(C)** Summary table of the proportion of dimerized particles in S proteins harbouring mutations at position 484. Asterisks indicate structures reported in previous publications^23^. **(D)** E484, K484, and Q484 RBD electrostatic surface potentials highlighting the surface potentials at position 484. The E484 surface potential was generated using the previously reported wild-type (D614G) + ACE2 focus-refined atomic model^23^. The K484 surface potential was generated using the previously reported D614G + N501Y + E484K focus-refined atomic model^23^. The Q484 surface potential was generated using the Kappa + ACE2 focus-refined atomic model reported in the present manuscript.

An analysis of the electrostatic surface potential at position 484 of the RBD reveals a unique property of the Kappa variant that may explain its propensity to dimerize. Figure 5D shows that the wild-type/Alpha/Delta variants and the Beta/Gamma variants may be binned into electronegative and electropositive surfaces at position 484, respectively, which would result in charge-charge repulsion if these variant S proteins were to dimerize in the same manner as the Kappa variant. The Kappa variant uniquely has an absence of charge at position 484 in its S protein, as reflected in the neutral surface potential shown in Figure 5D, consistent with its distinguishing ability to form head-to-head dimers.

## Discussion

Herein, we present a comparative analysis of the effects of the Delta and Kappa variant S protein mutations on aspects of viral fitness. Our overarching finding, as discussed below, is that *in the context of antibody evasion and ACE2 affinity, the Delta and Kappa variants don’t differ to a great degree, and that the global dominance of the Delta variant is likely a result of enhancements in other aspects of viral fitness*. Here, we discuss findings reported by others examining different aspects of viral fitness to rationalize the success of the Delta variant. Additionally, we highlight the Kappa variant S protein dimerization phenomenon as a distinguishing feature unique to this variant.

Antibody neutralization escape is an important element of increased viral fitness, with all previously characterized variants of concern exhibiting some degree of antibody escape^27,28,31, 32, 33^. Our antibody binding and neutralization experiments demonstrate that both variants escape monoclonal antibodies (via direct or potentially allosteric mutational mechanisms) and that these variants evade neutralization by vaccine-induced polyclonal antibody sera. These results are broadly consistent with initial reports on Kappa and Delta variant antibody escape^9,16,34,35^. Interestingly, a recent preprint - which includes a large panel of vaccine-induced sera samples - reports greater antibody evasion for the Kappa variant spike over the Delta variant^10^. There are several similar or identical mutations common between the Kappa/Delta variants and previously emerged variants of concern (Alpha, Beta, Gamma, Epsilon). For example, the L452R mutation, mutations at position 484 (E484K, E484Q), and mutations within the N3 loop in the NTD (residues 141-156) have all been previously characterized for their antibody-evasive effects in other variants. Our results demonstrating conserved mutational effects on antibody escape for these mutations in the Kappa and Delta variants - as seen in previously characterized variants of concern - further substantiates the modular nature of these mutations^23^.

We found unchanged and moderately increased ACE2 affinities for the Kappa and Delta variant spikes (compared to wild-type S protein) respectively. However, this moderate (∼2-fold) increase in ACE2 binding doesn’t rationalize the dominance of the Delta variant over other variants with higher fold-changes in ACE2 affinities (Alpha, Beta, Gamma)^27, 28, 29, 30, 31^. Analysis of the electrostatic surface potential of the ACE2-bound Delta variant S protein is consistent with enhancements in electrostatic complementarity afforded by both the L452R and T478K mutations. The same L452R mutation in the Kappa variant likely did not result in an overall change in ACE2 affinity due to the deleterious effects of a lost electrostatic interaction from the E484Q mutation. The only other study reporting Kappa/Delta S protein–ACE2 affinity differences (using RBDs alone) found effectively no change in affinity across ELISA, SPR, and BLI analyses^10^. Mutations outside of the RBD have however been demonstrated to influence ACE2 engagement by modulating RBD conformation, such as the D614G and A570D mutations in the B.1 and B.1.1.7 (Alpha variant) lineages respectively^4,25,36,37^. Therefore, the use of trimeric S proteins as employed in the present study may be important to realize allosteric effects on ACE2 binding difference by Kappa and Delta mutations outside of the RBD.

Biochemical analyses of the Kappa and Delta variant S proteins provide insights into the behaviour of the novel Delta plus (B.1.617.2 +) variant. The Delta plus variant contains the same mutations as the Delta variant, with the addition of a lysine to asparagine substitution at position 417 (K417N). The K417N mutation is found identically (K417N) and similarly (K417T) in the Beta (B.1.351) and Gamma (P.1) VoCs respectively ^1, 2, 3^. This mutation in isolation has previously been shown to enhance antibody evasion and concomitantly decrease ACE2 binding affinity^23^. The deleterious effect of decreased ACE2 affinity imparted by K417N/T mutation is compensated by N501Y substitution in Beta and Gamma VoCs, but not in Delta plus, potentially rationalizing its relatively limited global prevalence^38,39^. Consistent with previous mutational studies that used S proteins with the K417N mutation alone, recent results have demonstrated enhanced immune evasion and decreased ACE2 affinity for the Delta plus (B.1.617.2 +) variant relative to Delta (B.1.617.2)^10^. The consistency of the effects imparted by the K417N substitution, in isolation or when combined with other mutations, consolidates the understanding of naturally selected mutations as mostly independent functional modules at the molecular level. On this basis, the effects of rapidly emerging mutations that occur at a known location in the protein sequence can be accurately inferred from the in-depth structural and biophysical characterization of previous variants - even when these VoCs do not belong to the same lineage. Therefore, continued characterization of the effects of individual and combinations of mutations on overall viral fitness will allow researchers to better and more rapidly predict the behaviour of future emerging SARS-CoV-2 variants – with potential implications for vaccine modification and booster regimes.

The antibody evasion and ACE2 binding effects imparted by the Delta variant mutations are not unique to this VoC, yet the Delta variant has largely eclipsed other VoCs in terms of global prevalence (Figure 1D). Therefore, it is likely that mutational effects that influence other aspects of viral fitness (not assessed in the present publication) further rationalize the unique current prevalence of the Delta lineage. For example, the P681R mutation - found in all B.1.617 sub-lineages - has recently been reported to enhance S protein cleavage and cell-cell fusion^11,15,35^. As S protein cleavage is a crucial step in the pre- to post-fusion transition of the S protein, the P681R mutation likely contributes to the reported increase in Delta variant replication kinetics and viral loads in oropharyngeal and nose/throat swabs of infected individuals^16,40^. However, given the shared P681R mutation in both Kappa and Delta variants, this mutation alone does not explain the prevalence of the Delta variant over Kappa in India initially and subsequently globally. Finally, the effects of the D950N mutation (Delta) and the Q1071H mutation within the S protein fusion machinery (Kappa), as well as mutations outside of the S protein open-reading frame, have yet to be assessed for their impact on SARS-CoV-2 viral fitness.

Consistent with published structures of other SARS-CoV-2 VoC S proteins, our structure of the Delta S protein revealed no large changes in global 3D structure (Figure S11). However, for the Kappa variant we report an unprecedented head-to-head S trimer dimerization, likely facilitated by the abrogation of charge-charge repulsion and the additional contacts afforded by the E484Q substitution. Our synthetic mutation of Q484I in the Kappa variant background, which also resulted in dimerization, suggests that the SARS-CoV-2 S protein may be a single amino acid substitution away from exhibiting this dimer-of-trimers phenotype. However, an analysis of mutational frequency at position 484 in globally deposited sequences reveals that only E484, K484, and Q484 S protein genotypes have ever been present at >1% of total deposited sequences, suggesting limited mutational flexibility at this position (Figure S12). We found this head-to-head dimerization to be concentration-dependent, with no evidence of dimerization in experiments conducted at <0.05 mg/mL (see size-exclusion chromatography, Figure S1, and negative-stain electron microscopy, Figure S10). Thus, we hypothesize that if local spike concentrations reach high enough concentrations to dimerize at any point during the SARS-CoV-2 cell entry, replication, and packaging events, that this dimerization phenomenon could have biological implications.

While future studies will be required to assess the potential biological relevance of the reported dimerization of spike trimers, we speculate here on some mechanisms by which dimerized S proteins could result in increased or decreased viral fitness. Firstly, head-to-head S protein dimerization buries much of the antibody-accessible surface area of the RBD (the predominant target of neutralizing antibodies) and could shield this otherwise vulnerable neutralization site^13^. Secondly, dimerized spikes - in the same manner as reported for the Kappa variant - would be unable to engage the ACE2 receptor and therefore not be able to enter host cells through the ACE2-dependent cell-entry pathway^41^. These first two competing mechanisms resulting in increased and decreased viral fitness, respectively, could favour a spike protein with a finely tuned balance of dimerization potential to both mask neutralizing epitopes, but also to readily dissociate and permit engagement of ACE2. To this second point, we verified that the Kappa S protein dimer-of-trimers complex is labile enough to still permit ACE2 binding through our experimental derivation of the ACE2 bound structure. We saw no evidence of S protein dimer-of-trimer formation in our cryo-EM images upon introducing a modest excess of ACE2 (∼1:1.25 S protein trimer : ACE2 molar ratio), despite an identical S protein concentration which resulted in the dimer-of-trimers reconstruction. Interestingly, two recent publications have independently described potent neutralizing nanobodies with propensities to induce S protein dimers^42,43^. While the exact mode of neutralization for these nanobodies remains unclear, this may suggest that S protein dimerization has negative impacts on viral fitness. Further studies to elucidate the biological implications, if any, of this dimerization phenomenon are therefore highly relevant in the rapidly evolving SARS-CoV-2 variant landscape.

## Acknowledgments

This work was supported by awards to S.S. from a Canada Excellence Research Chair Award, the VGH Foundation, Genome BC, Canada, and from the Tai Hung Fai Charitable Foundation. W.L. and D.S.D. were supported by the UPMC. J.W.S is supported by a CIHR Frederick Banting and Charles Best Canada Graduate Scholarships Doctoral Award (CGS D) and a UBC President’s Academic Excellence Initiative PhD Award. D.M. is supported by a CIHR Frederick Banting and Charles Best Canada Graduate Scholarship Master’s Award (CGS-M). J.-P.D. is supported by a Long-Term Fellowship from the Human Frontier Science Program.

## Author Contributions

J.W.S., D.M., S.Z. carried out expression and purification of the spike proteins and antibodies. J.W.S and D.M. performed the molecular cloning. A.K. performed the BLI binding assays under supervision from W.L. and D.S.D.. J.W.S. and D.M. performed the antibody binding experiments. D.M. performed the antibody neutralization experiments. I.S. provided the vaccine-induced patient-derived sera samples and aided the interpretation of the data. A.M.B., J.P.D., and K.S.T. carried out the experimental components of cryo-EM and electron microscopy including specimen preparation and data collection. X.Z. carried out all computational aspects of image processing and structure determination. D.M., J.W.S., X.Z., S.S.S., and S.S. interpreted and analyzed the cryo-EM structures. A.K., W.L., and D.S.D. provided the plasmids for VH-ab8 and IgG ab1 as part of a collaboration between the Subramaniam and Dimitrov laboratories on SARS-CoV-2. J.W.S., D.M., and S.S. drafted the initial manuscript with input from all authors.

## Materials and Methods

### Kits and reagents

|  |  |  |
| --- | --- | --- |
| Pierce 1-Step Ultra Substrate Solution | Thermo Fisher Scientific | Cat. # 34028 |
| Q5 Site-Directed mutagenesis kit | NEB | Cat. # E0554S |
| Lenti-X™ GoStix™ Plus | Takara Bio | Cat. # 631281 |
| ONE-Glo™ EX Luciferase Assay System | Promega | Cat. # E8130 |

### Antibodies

|  |  |  |
| --- | --- | --- |
| VH ab8 | 17, 18 | N/A |
| IgG1 ab1 | 17, 18 | N/A |
| Fab S309 | 22 | N/A |
| Fab S2M11 | 21 | N/A |
| Fab 4A8 | 19 | N/A |
| Fab 4-8 | 20 | N/A |
| goat anti-human IgG | Jackson ImmunoResearch | Cat. # 109-035-088 |

### Recombinant proteins

|  |  |  |
| --- | --- | --- |
| Ace2 (18-615) | New England Biolabs | Cat. # 73775S |

### Cell lines

|  |  |  |
| --- | --- | --- |
| Expi293F | Thermo Fisher Scientific | Cat# A14527 |

### Recombinant DNA and Oligonucleotides

|  |  |  |
| --- | --- | --- |
| pαH SARS-CoV-2 S HexaPro | 44 | Addgene<br>#154754 |
| pαH HexaPro D614G | 23 | N/A |
| pcDNA3.1 Fab S309 Light Chain | 23 | N/A |
| pcDNA3.1 Fab S309 Heavy Chain | 23 | N/A |
| pcDNA3.1 Fab S2M11 Light Chain | 23 | N/A |
| pcDNA3.1 Fab S2M11 Heavy Chain | <sup>23</sup> | N/A |
| pcDNA3.1 Fab 4A8 Light Chain | This paper | N/A |
| pcDNA3.1 Fab 4A8 Heavy Chain | This paper | N/A |
| pcDNA3.1 Fab 4-8 Light Chain | This paper | N/A |
| pcDNA3.1 Fab 4-8 Heavy Chain | This paper | N/A |

### Software and Algorithms

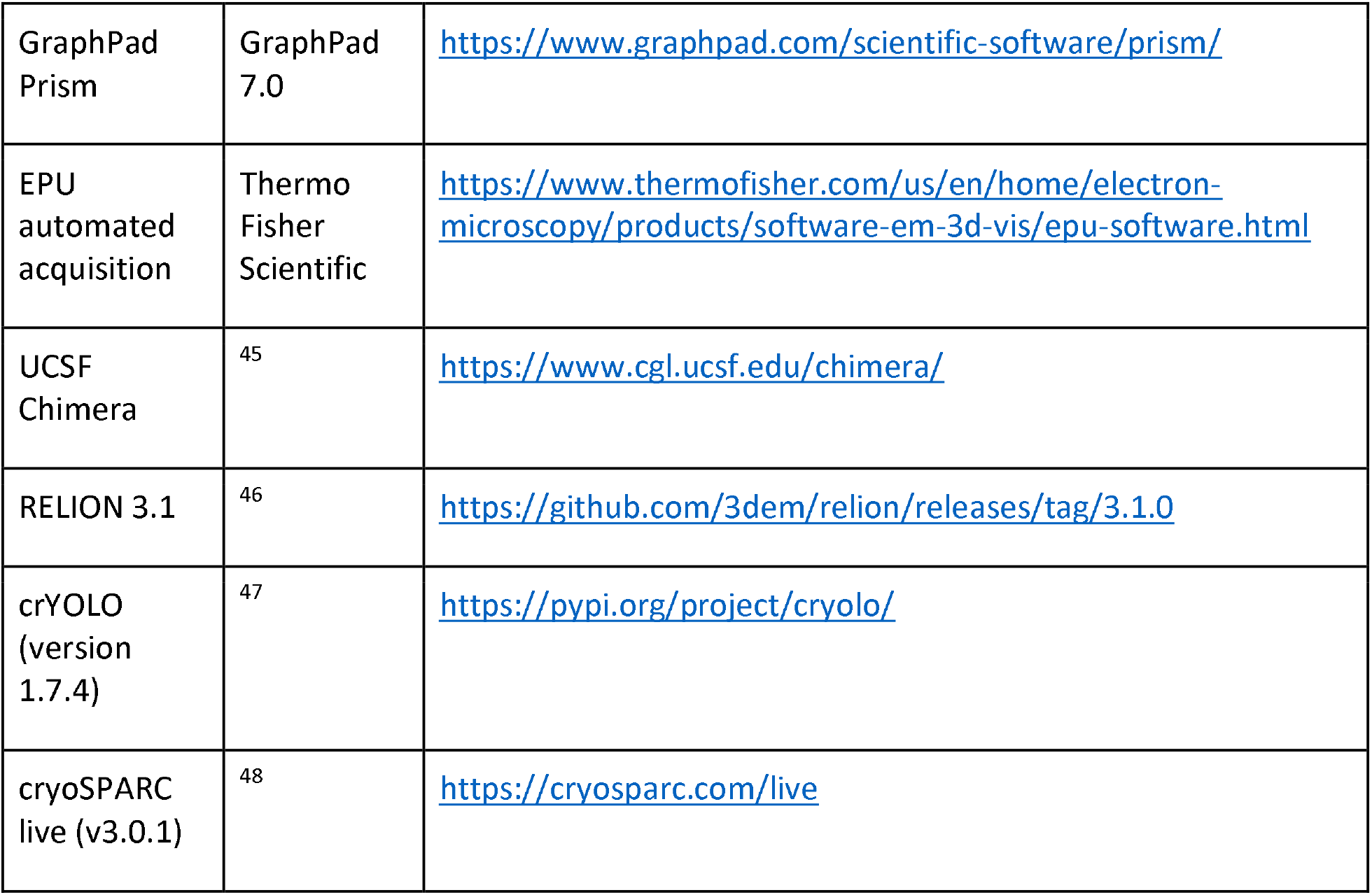

## Method details

### Pseudovirus Neutralization Assay

SARS-CoV-2 S protein Delta and Kappa genes were synthesized and inserted into pcDNA3.1 (GeneArt Gene Synthesis, Thermo Fisher Scientific). The production of the SARS-CoV-2 wild-type (D614G) S protein was described previously^23^. Variant pseudotyped retroviral particles were produced in HEK293T cells as described previously^49^. Briefly, a third-generation lentiviral packaging system was utilized in combination with plasmids encoding the full-length SARS-CoV-2 spike, along with a transfer plasmid encoding luciferase and GFP as a dual reporter gene. Pseudoviruses were harvested 60 h after transfection, filtered with a 0.45 µm PES filter, and frozen. For neutralization assays, HEK293T-ACE2-TMPRSS2 cells^50^ (BEI Resources cat# NR-55293) were seeded in 96- or 384-well plates at 50 000 or 20 000 cells respectively. The next day, pseudovirus preparations normalized for viral capsid p24 levels (Lenti-X™ GoStix™ Plus) were incubated with dilutions of the indicated antibodies or sera, or media alone for 1 h at 37°C prior to addition to cells and incubation for 48 h. Cells were then lysed and luciferase activity assessed using the ONE-Glo™ EX Luciferase Assay System (Promega) according to the manufacturer’s specifications. Detection of relative luciferase units was carried out using a Varioskan Lux plate reader (Thermo Fisher). Percent neutralization was calculated relative to signals obtained in the presence of virus alone for each experiment.

### Expression and Purification of Recombinant Spike Protein Constructs

The wild-type SARS-CoV-2 S HexaPro expression plasmid was previously described^44^ and was a gift from Jason McLellan (Addgene plasmid #154754; http://n2t.net/addgene:154754; RRID:Addgene_154754).

The VoC RBD mutations were introduced by site-directed mutagenesis (Q5 Site-Directed Mutagenesis Kit, New England Biolabs). Successful cloning was confirmed by Sanger sequencing (Genewiz, Inc.). Expi293F cells (Thermo Fisher, Cat# A14527) were grown in suspension culture using Expi293 Expression Medium (Thermo Fisher, Cat# A1435102) at 37°C, 8% CO_2_. Cells were transiently transfected at a density of 3 × 10^6 cells/mL using linear polyethylenimine (Polysciences Cat# 23966-1). The media was supplemented 24 hours after transfection with 2.2 mM valproic acid, and expression was carried out for 3–5 days at 37°C, 8% CO_2_. The supernatant was harvested by centrifugation and filtered through a 0.22-μM filter prior to loading onto a 5 mL HisTrap excel column (Cytiva). The column was washed for 20 CVs with wash buffer (20 mM Tris pH 8.0, 500 mM NaCl), 5 CVs of wash buffer supplemented with 20 mM imidazole, and the protein eluted with elution buffer (20 mM Tris pH 8.0, 500 mM NaCl, 500 mM imidazole). Elution fractions containing the protein were pooled and concentrated (Amicon Ultra 100 kDa cut off, Millipore Sigma) for gel filtration. Gel filtration was conducted using a Superose 6 10/300 GL column (Cytiva) pre-equilibrated with GF buffer (20 mM Tris pH 8.0, 150 mM NaCl). Peak fractions corresponding to soluble protein were pooled and concentrated to 4.5–5.5 mg/mL (Amicon Ultra 100 kDa cut off, Millipore Sigma). Protein samples were flash-frozen in liquid nitrogen and stored at -80**°**C.

Antibody Production

VH-FC ab8, IgG ab1, Fab S309, and Fab S2M11 were produced as previously described^17, 18^. Plasmids encoding light and heavy chains for Fab 4A8 and Fab 4-8 were synthesized (GeneArt Gene Synthesis, Thermo Fischer Scientific). Heavy chains were designed to incorporate a C terminal 6x histidine tag. Expi293 cells were transfected at a density of 3 × 10^6 cells/mL using linear polyethylenimine (Polysciences Cat# 23966-1). 24-hours following transfection, media was supplemented with 2.2 mM valproic acid, and expression was carried out for 3–5 days at 37°C, 8% CO_2_. The supernatant was harvested by centrifugation and filtered through a 0.22 μM filter prior to loading onto a 5 mL HisTrap excel column (Cytiva). The column was washed for 20 CVs with wash buffer (20 mM Tris pH 8.0, 500 mM NaCl), 5 CVs of wash buffer supplemented with 20 mM imidazole. The protein was eluted with elution buffer (20 mM Tris pH 8.0, 500 mM NaCl, 500 mM imidazole). Elution fractions containing the protein were pooled and concentrated (Amicon Ultra 10 kDa cut off, Millipore Sigma) for gel filtration. Gel filtration was conducted using a Superose 6 10/300 GL column (Cytiva) pre-equilibrated with GF buffer (20 mM Tris pH 8.0, 150 mM NaCl). Peak fractions corresponding to soluble protein were pooled and concentrated to 8–20 mg/mL (Amicon Ultra 10 kDa cut off, Millipore Sigma). Protein samples were stored at 4°C until use.

### Electron Microscopy Sample Preparation and Data Collection

For cryo-EM, S protein samples were prepared at 2.25 mg/mL, with and without the addition of ACE2 at 0.5 mg/mL (1:1.25 S protein trimer:ACE2 molar ratio). Vitrified samples of all S protein samples were prepared by first glow discharging Quantifoil R1.2/1.3 Cu mesh 200 holey carbon grids for 30 seconds using a Pelco easiGlow glow discharge unit (Ted Pella) and then applying 1.8 µL of protein suspension to the surface of the grid at a temperature of 10°C and a humidity level of >98%. Grids were blotted (12 sec, blot force -10) and plunge frozen into liquid ethane using a Vitrobot Mark IV (Thermo Fisher Scientific). All cryo-EM samples were imaged using a 300 kV Titan Krios G4 transmission electron microscope (Thermo Fisher Scientific) equipped with a Falcon4 direct electron detector in electron event registration (EER) mode. Movies were collected at 155,000x magnification (calibrated pixel size of 0.5 Å per physical pixel) over a defocus range of -0.5 µm to -3 µm with a total dose of 40 e^-^/Å^2^ using EPU automated acquisition software. For negative stain, S protein samples were prepared at 0.1 mg/mL. Grids of Cu mesh 300 with continuous ultra-thin carbon film (CF300-Cu-UL, Electron Microscopy Sciences) were glow discharged for 15 seconds using the Pelco easiGlow. Samples were allowed to adsorb for 30 seconds before blotting away excess liquid, followed by a brief wash with MilliQ H_2_O. Grids were stained by three successive applications of 2% (w/v) uranyl formate (20 s, 20 s, 60 s). Negative stain grids were imaged using a 200 kV Glacios transmission electron microscope (Thermo Fisher Scientific) equipped with a Falcon3 camera operated in linear mode. Micrographs were collected using EPU at 92,000x magnification (physical pixel size 1.6 Å) over a defocus range of -1 µm to -2 µm with a total accumulated dose of 120 e^-^/Å^2^.

### Image Processing

The detailed workflow for the data processing is summarized in Supplementary Figures S3-S9. In general, all data processing was performed in cryoSPARC v.3.2^48^ unless stated otherwise. Motion correction in patch mode (EER upsampling factor 1, EER number of fractions 40), CTF estimation in patch mode, reference-free particle picking, and particle extraction were performed on-the-fly in cryoSPARC. After preprocessing, particles were subjected to 2D classification and/or 3D heterogeneous classification. The final 3D refinement included per particle CTF estimation and aberration correction. For B.1.617.1 spike proteins, focused refinements were performed with a soft mask covering all six RBDs. For the complexes of spike protein ectodomain and human ACE2, focused refinements were performed with a soft mask covering a single RBD and its bound ACE2. Global resolution and focused resolution were determined according to the gold-standard FSC^51^.

### Model Building and Refinement

For models of spike protein ectodomain alone, the SARS-CoV-2 HexaPro S trimer with N501Y mutation (PDB code 7MJG) was docked into the cryo-EM density map using UCSF Chimera v.1.15^45^. Then, mutation and manual adjustment were done with COOT v.0.9.3^52^, followed by iterative rounds of refinement in COOT and Phenix v.1.19^53^. Glycans were added at N-linked glycosylation sites in COOT. For models of complex of spike protein ectodomain and human ACE2, the RBD-ACE2 subcomplex was built using published coordinates (PDB code 7MJN) as the initial model, followed by refinement against focused refinement maps. The obtained model was then docked into global refinement maps together with the other individual domains of the spike protein. Model validation was performed using MolProbity^54^. Figures were prepared using UCSF Chimera, UCSF ChimeraX v.1.1.1^55^, and PyMOL (v.2.2 Schrodinger, LLC).

### Biolayer Interferometry (BLI) S protein-ACE2 Binding Assay

The kinetics of SARS-CoV-2 trimers and human ACE2 binding were analyzed with the biolayer interferometer BLItz (ForteBio, Menlo Park, CA). Protein-A biosensors (ForteBio: 18–5010) were coated with ACE2-mFc (40 µg/mL) for 2 min and incubated in DPBS (pH = 7.4) to establish baselines. Concentrations of 125, 250, 500, and 1000 nM spike trimers were used for association for 2 min followed by dissociation in DPBS for 5 min. The association (*k*_*on*_) and dissociation (*k*_*off*_) rates were derived from the fitting of sensorgrams and used to calculate the binding equilibrium constant (K_D_).

### Enzyme-Linked Immunosorbent Assay (ELISA)

For ELISA, 100 µL of wild-type (D614G), Kappa, or Delta SARS-CoV-2 S proteins were coated onto 96-well MaxiSorp™ plates at 2 µg/mL in PBS + 1% casein overnight at 4°C. All washing steps were performed 3 times with PBS + 0.05% Tween 20 (PBS-T). After washing, wells were incubated with blocking buffer (PBS-T + 1% casein) for 1 hr at room temperature. After washing, wells were incubated with dilutions of primary antibodies in PBS-T + 0.5% BSA buffer for 1 hr at room temperature. After washing, wells were incubated with goat anti-human IgG (Jackson ImmunoResearch) at a 1:8,000 dilution in PBS-T + 1% casein buffer for 1 hr at room temperature. After washing, the substrate solution (Pierce™ 1-Step™) was used for colour development according to the manufacturer’s specifications. Optical density at 450 nm was read on a Varioskan Lux plate reader (Thermo Fisher Scientific).

