## Supplemental Information for "Structural and Biochemical Rationale for Enhanced Spike Protein Fitness in Delta and Kappa SARS-CoV-2 Variants"

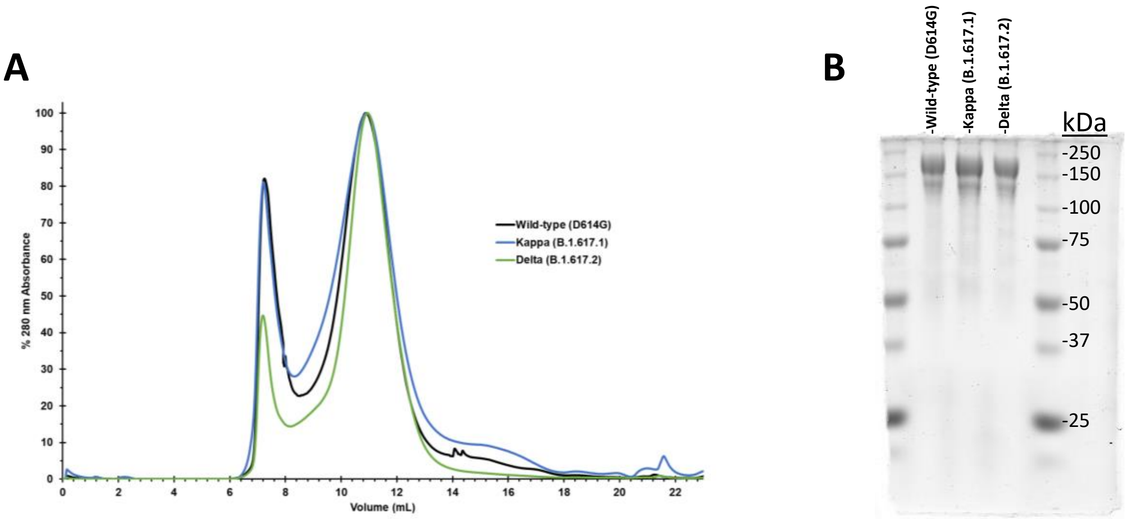


**Figure S1: Purification of wild-type (D614G), Kappa (B.1.617.1), and Delta (B.1.617.2) spike protein ectodomains. (A)** Superose 6 10/300 GL size-exclusion traces for the spike protein ectodomains employed in this study. **(B)** SDS polyacrylamide gel electrophoresis (SDS-PAGE) gel of the three SARS-CoV-2 S proteins employed in this study. A normalized amount (4 µg) of each variant spike protein was loaded in each well.

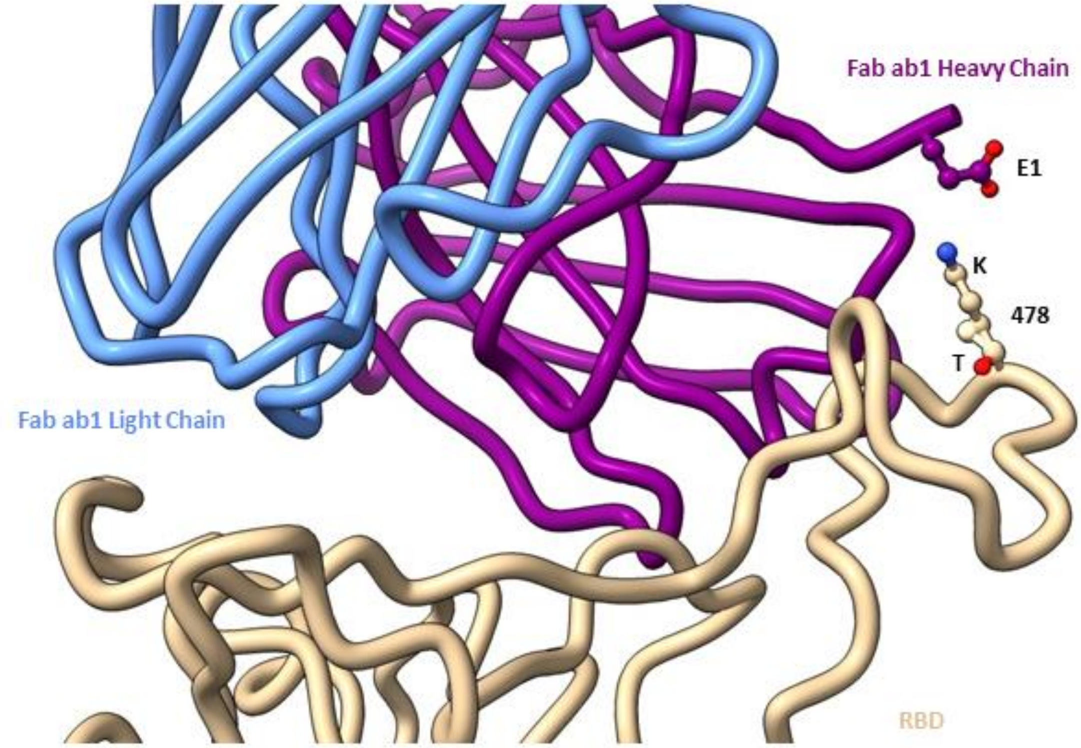


**Figure S2: Proximity of position 478 within the Delta (B.1.617.2) Variant RBD to Glutamic acid 1 within Fab ab1.** The model resulting from focused refinement of the interface between the N501Y mutant spike and Fab ab1 was used (PDB: 7MJL). The proposed positioning of K478 is shown overlapping the experimental positioning of T478.

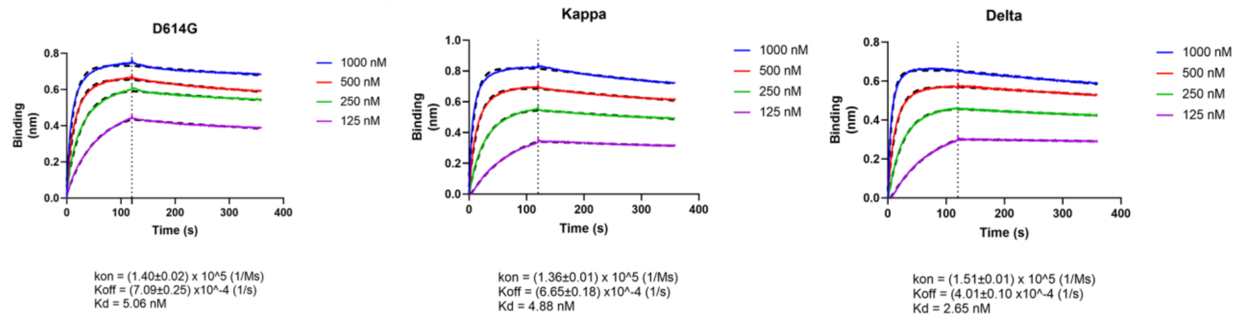


**Figure S3: Raw biolayer interferometry (BLI) sensorgrams for ACE2 binding to wild-type (D614G), Kappa (B.1.617.1), and Delta (B.1.617.2) spike protein ectodomains.** ACE2 was immobilized on BLI sensor tips and S proteins were assessed for binding at different concentrations as indicated. Biophysical parameters ($K_D$, $k_{on}$, $k_{off}$) are shown as means. Errors correspond to standard deviations.

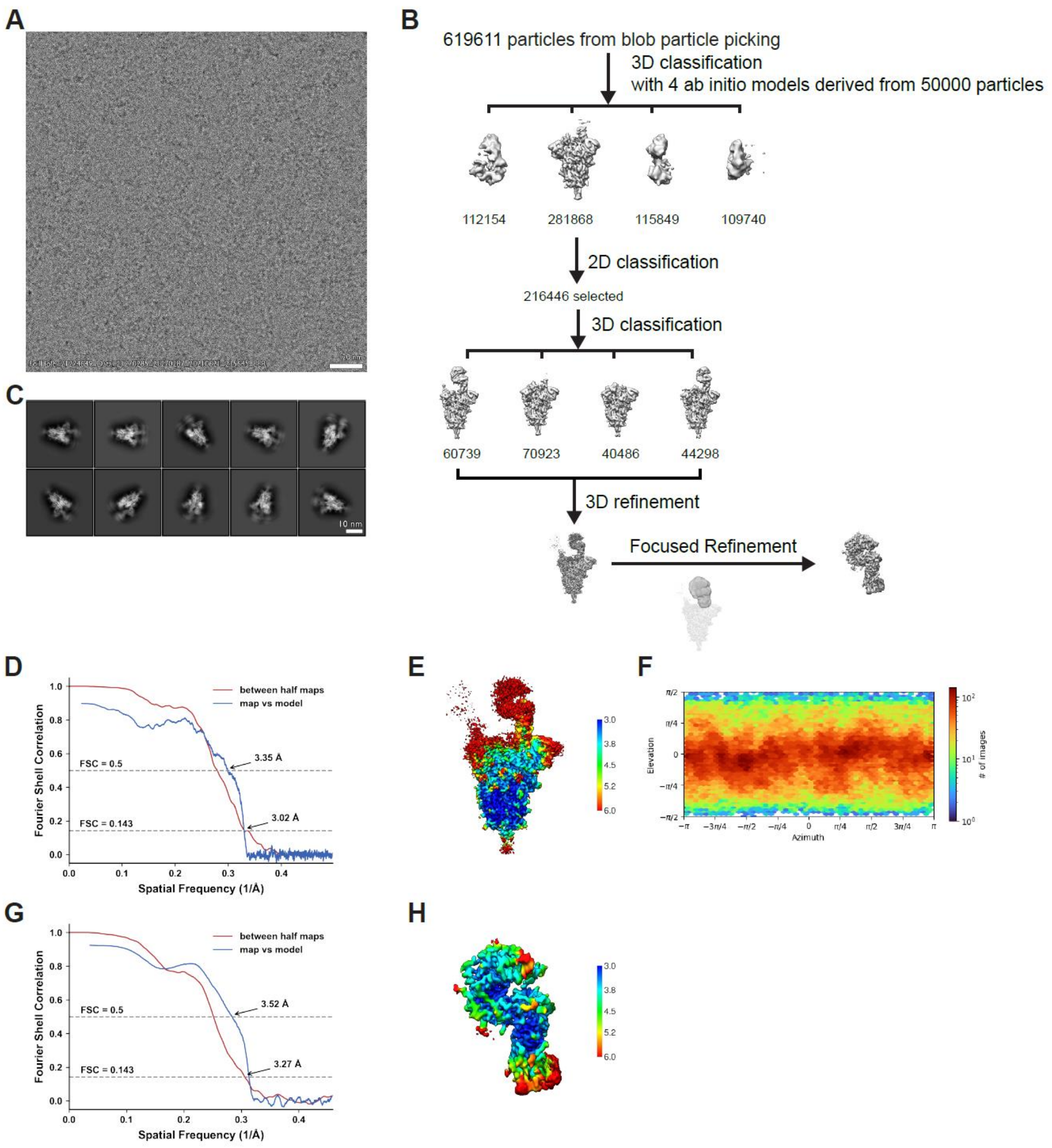


**Figure S4. Cryo-EM data processing and validation for the Kappa (B.1.617.1) spike protein ectodomain in complex with ACE2. (A)** Representative cryo-EM micrograph. **(B)** Workflow of cryo-EM image processing. **(C)** Representative 2D classes. **(D)** Fourier Shall Correlation (FSC) curves for the global refinement. **(E)** Local resolution map for the global refinement. **(F)** Viewing direction distribution plot. **(G)** Fourier Shall Correlation (FSC) curves for the focused refinement of the RBD-ACE2 interface. **(H)** Local resolution map for the focused refinement.

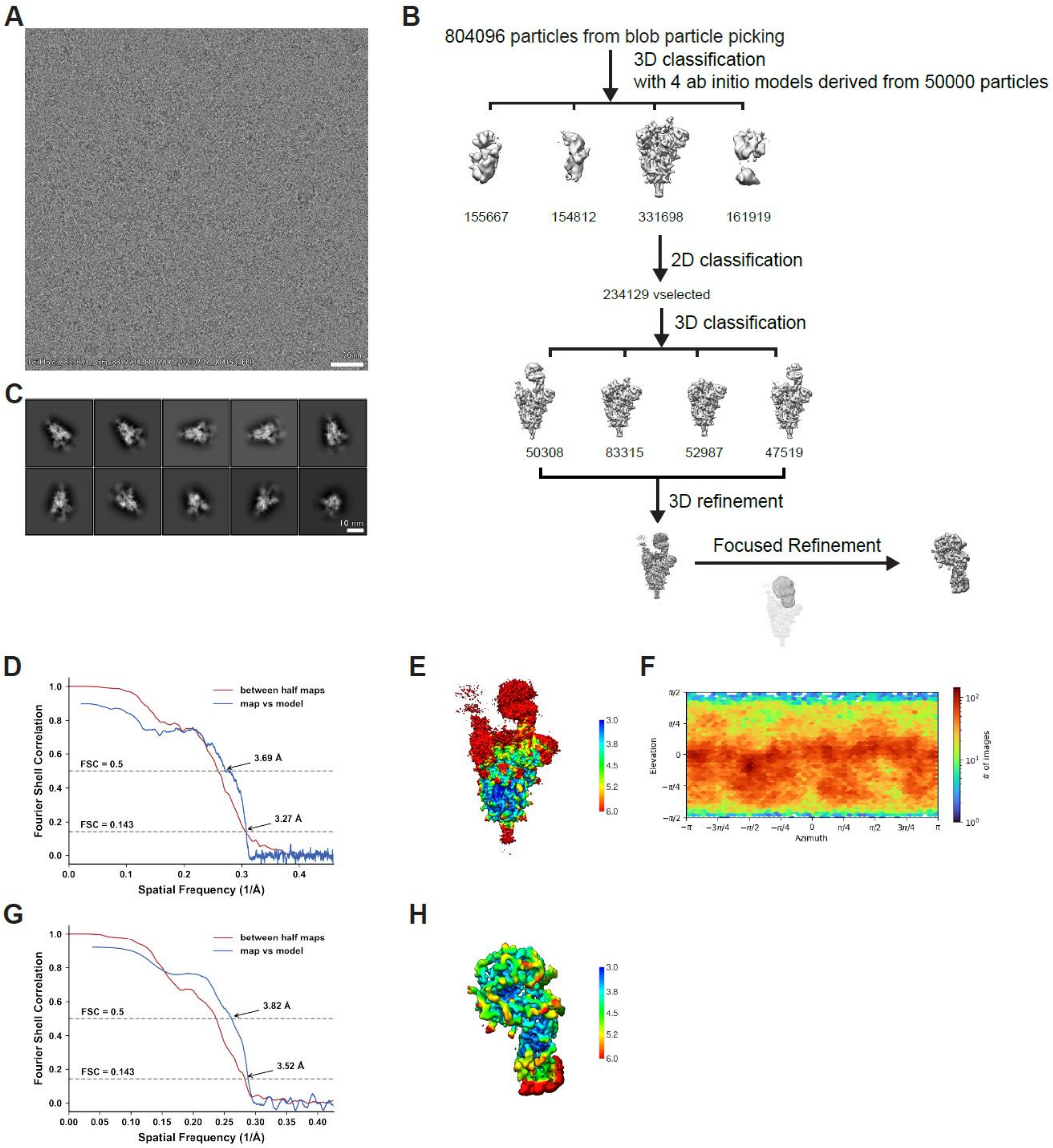


**Figure S5. Cryo-EM data processing and validation for the Delta (B.1.617.2) spike protein ectodomain in complex with ACE2. (A)** Representative cryo-EM micrograph. **(B)** Workflow of cryo-EM image processing. **(C)** Representative 2D classes. **(D)** Fourier Shall Correlation (FSC) curves for the global refinement. **(E)** Local resolution map for the global refinement. **(F)** Viewing direction distribution plot. **(G)** Fourier Shall Correlation (FSC) curves for the focused refinement of the RBD-ACE2 interface. **(H)** Local resolution map for the focused refinement.

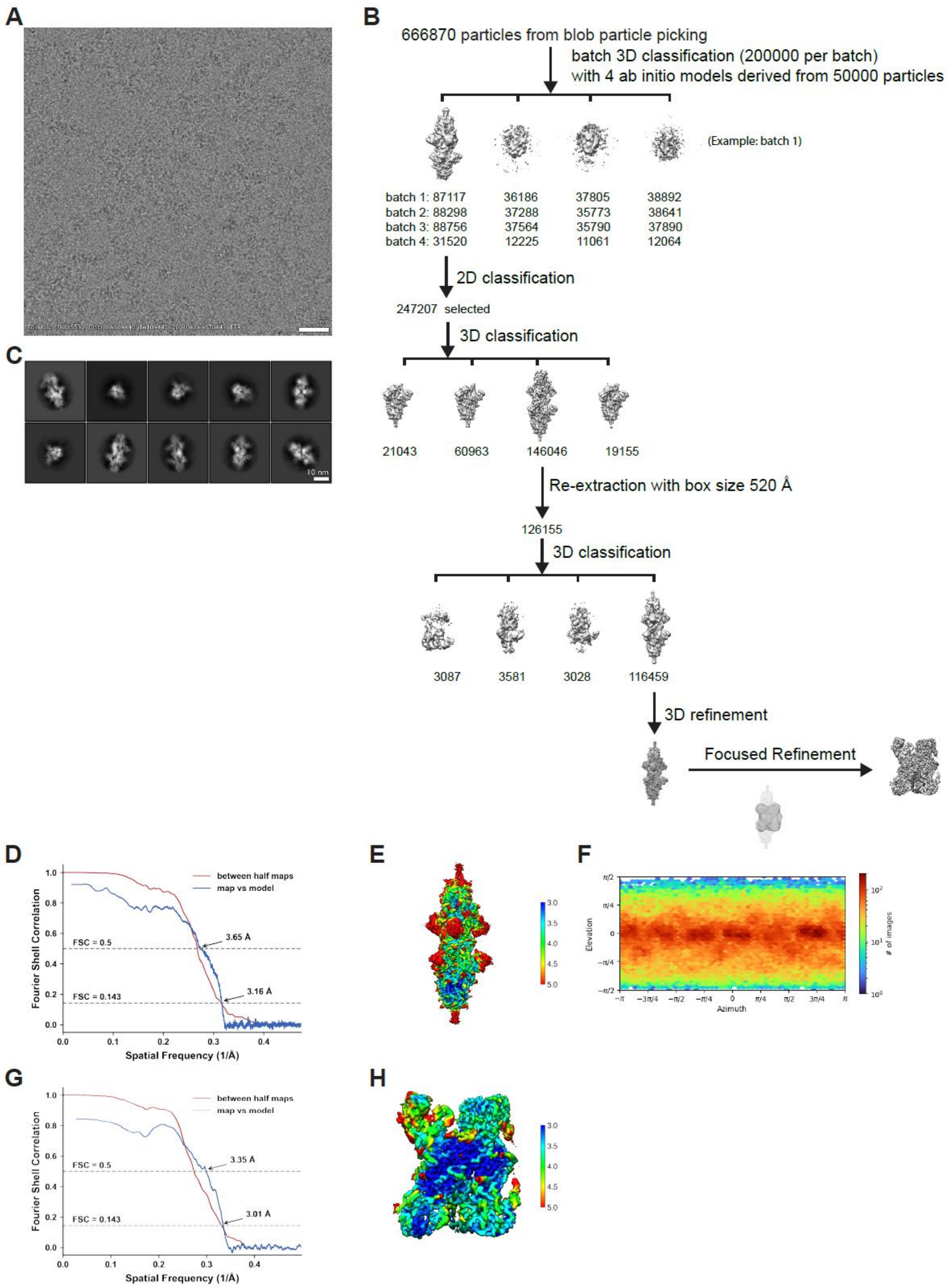


**Figure S6. Cryo-EM data processing and validation for the Kappa (B.1.617.1) spike protein ectodomain.** **(A)** Representative cryo-EM micrograph. **(B)** Workflow of cryo-EM image processing. **(C)** Representative 2D classes. **(D)** Fourier Shall Correlation (FSC) curves for the global refinement. **(E)** Local resolution map for the global refinement. **(F)** Viewing direction distribution plot. **(G)** Fourier Shall Correlation (FSC) curves for the focused refinement of the dimer interface. **(H)** Local resolution map for the focused refinement.

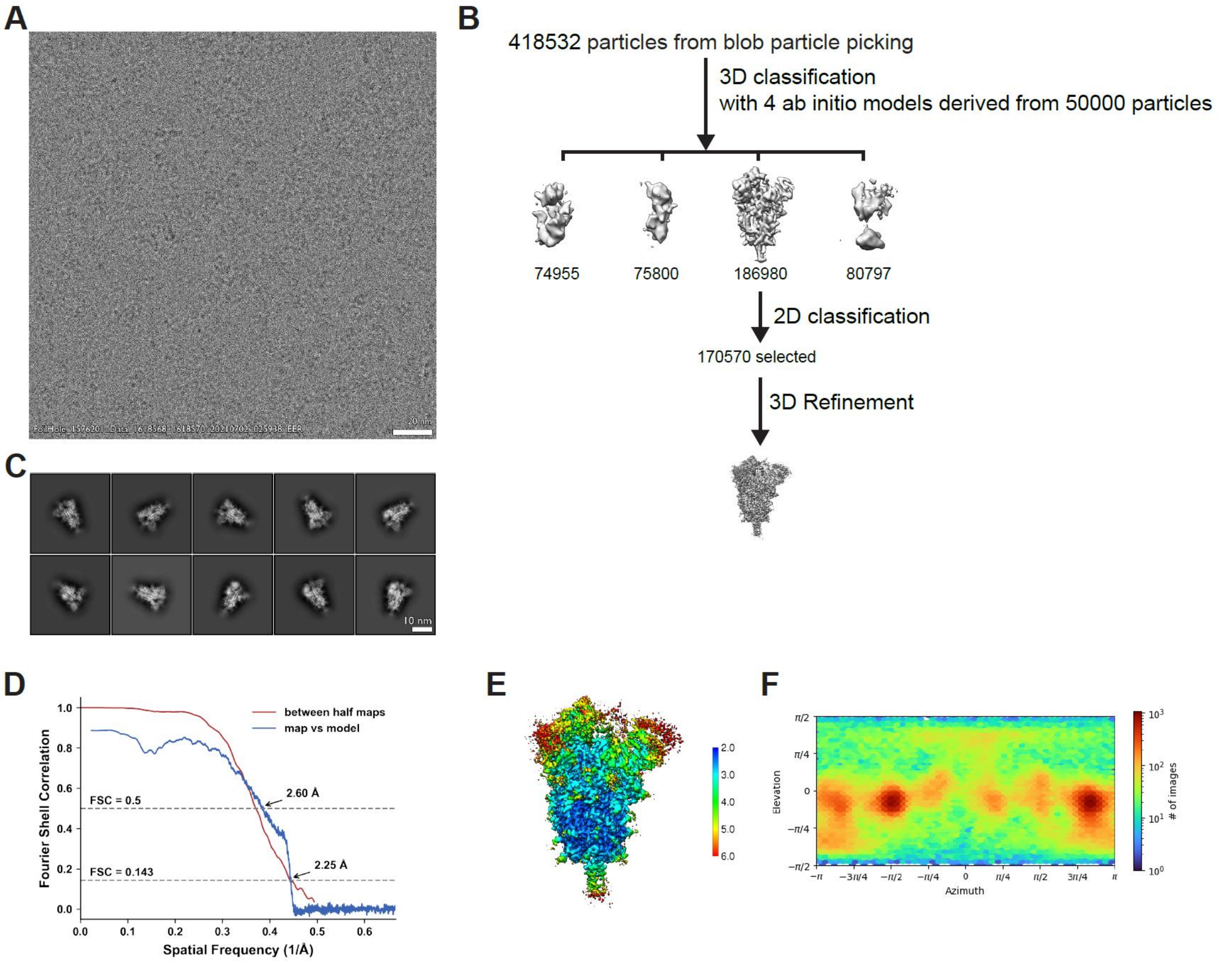


**Figure S7. Cryo-EM data processing and validation for the Delta (B.1.617.2) spike protein ectodomain.** **(A)** Representative cryo-EM micrograph. **(B)** Workflow of cryo-EM image processing. **(C)** Representative 2D classes. **(D)** Fourier Shall Correlation (FSC) curves for the global refinement. **(E)** Local resolution map for the global refinement. **(F)** Viewing direction distribution plot.

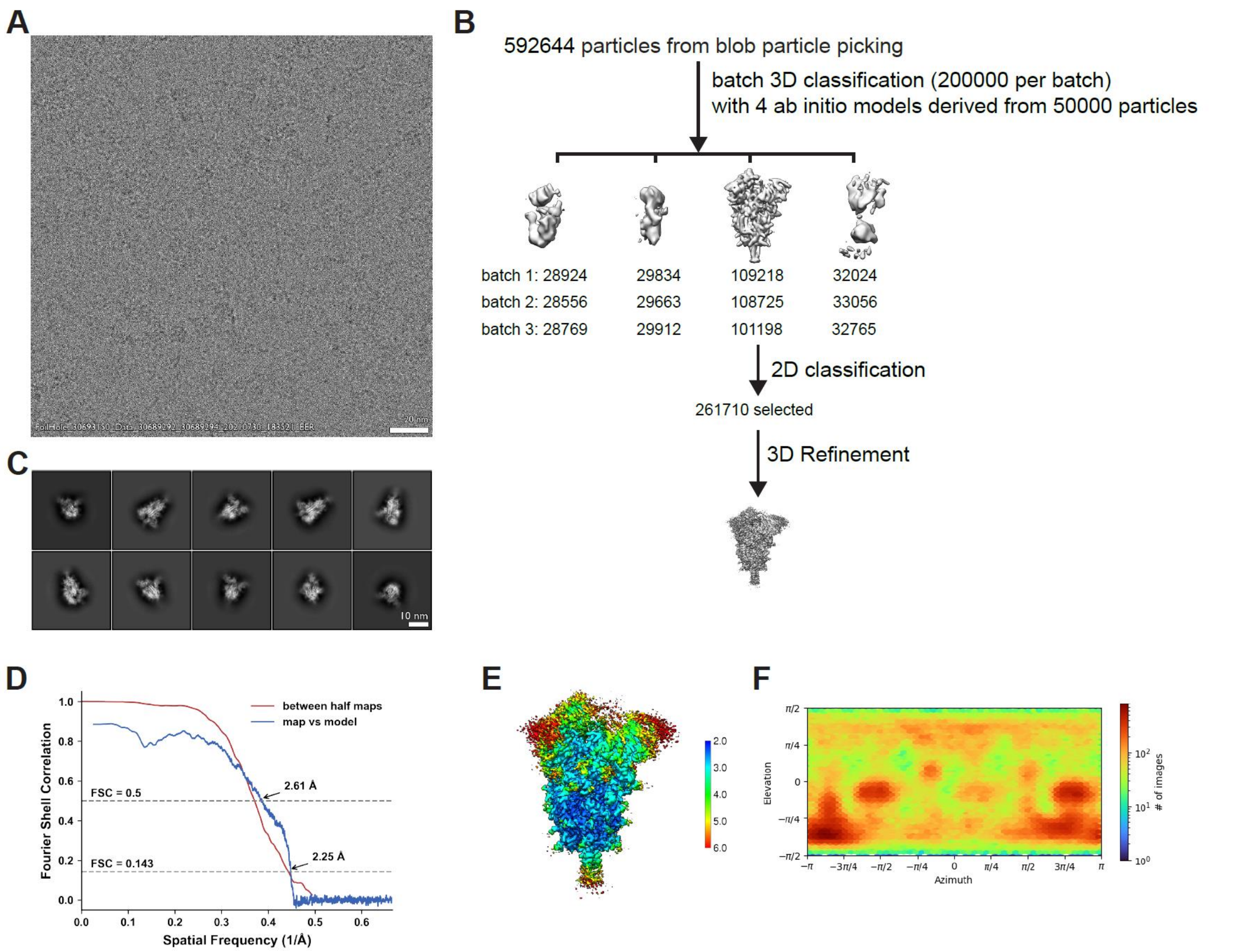


**Figure S8. Cryo-EM data processing and validation for the Kappa (B.1.617.1) + Q484A spike protein ectodomain. (A)** Representative cryo-EM micrograph. **(B)** Workflow of cryo-EM image processing. **(C)** Representative 2D classes. **(D)** Fourier Shall Correlation (FSC) curves for the global refinement. **(E)** Local resolution map for the global refinement. **(F)** Viewing direction distribution plot.

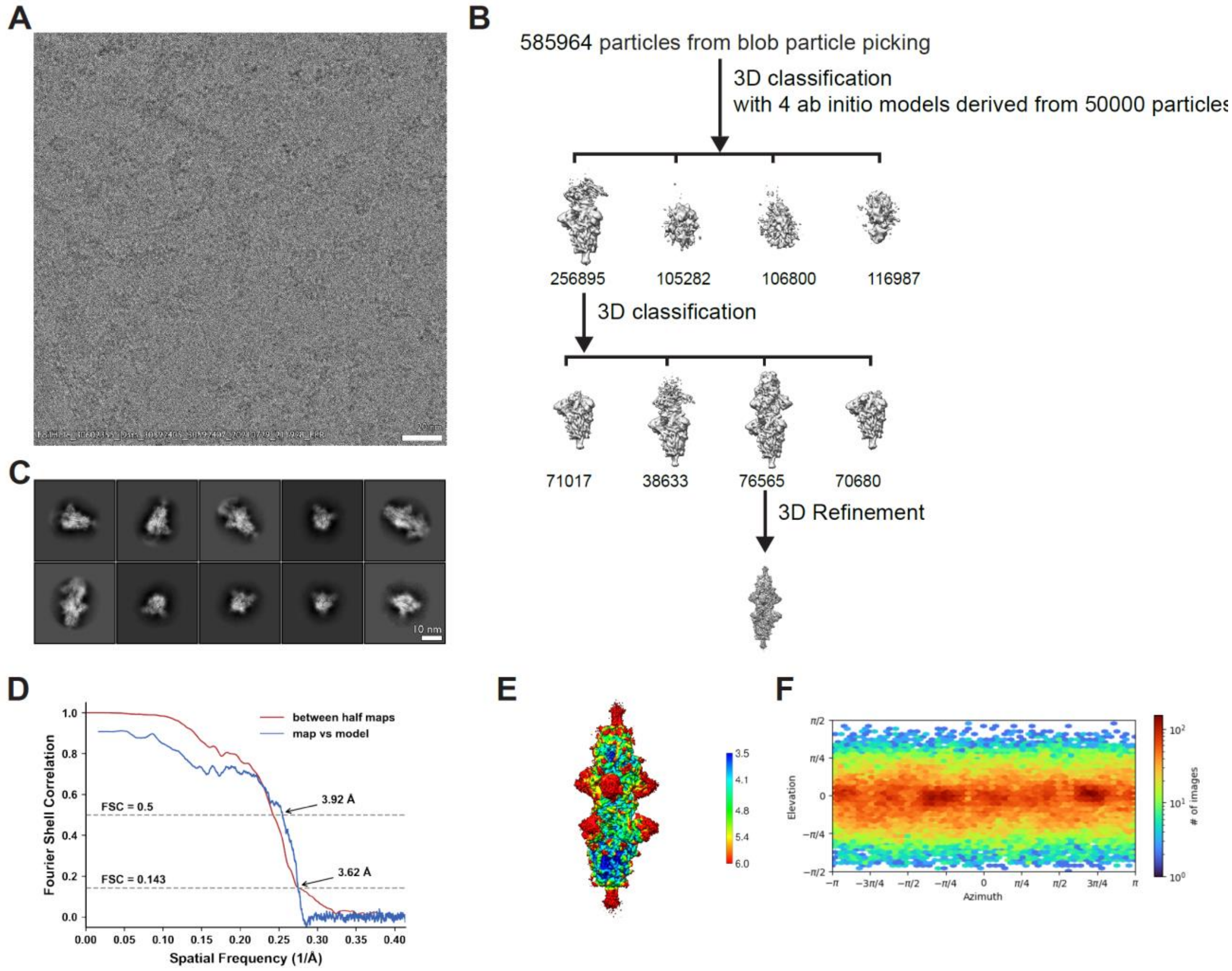


**Figure S9. Cryo-EM data processing and validation for the Kappa (B.1.617.1) + Q484I spike protein ectodomain. (A)** Representative cryo-EM micrograph. **(B)** Workflow of cryo-EM image processing. **(C)** Representative 2D classes. **(D)** Fourier Shall Correlation (FSC) curves for the global refinement. **(E)** Local resolution map for the global refinement. **(F)** Viewing direction distribution plot.

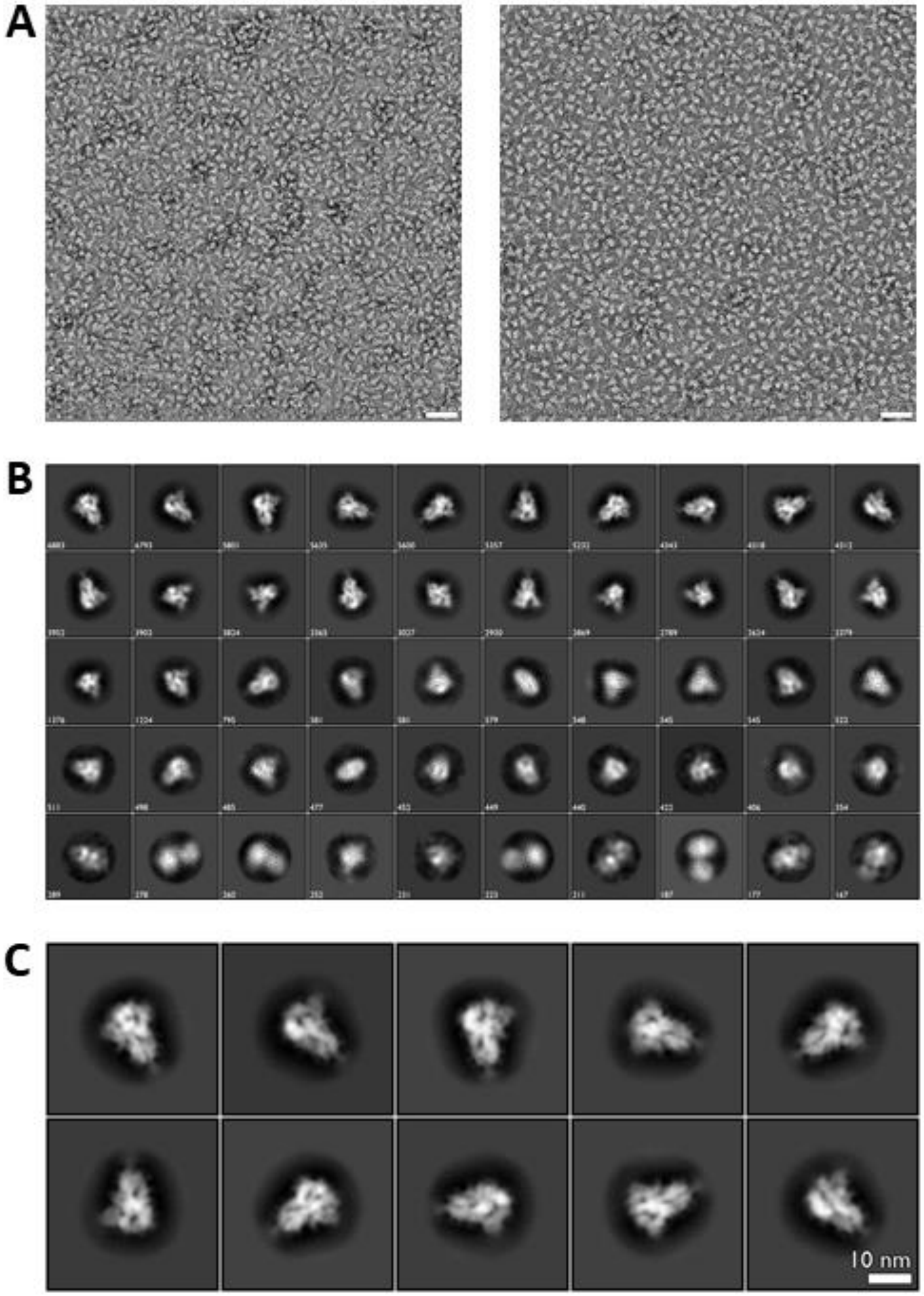


**Figure S10: The Kappa (B.1.617.1) variant S protein ectodomain reveals no dimerization under negative stain electron microscopy conditions. (A)** Two representative micrographs selected from the total dataset for the Kappa (B.1.617.1) variant S protein ectodomain. **(B)** All 2D class averages derived from the micrographs as represented in (A). **(C)** The ten most highly populated 2D class averages from (B).

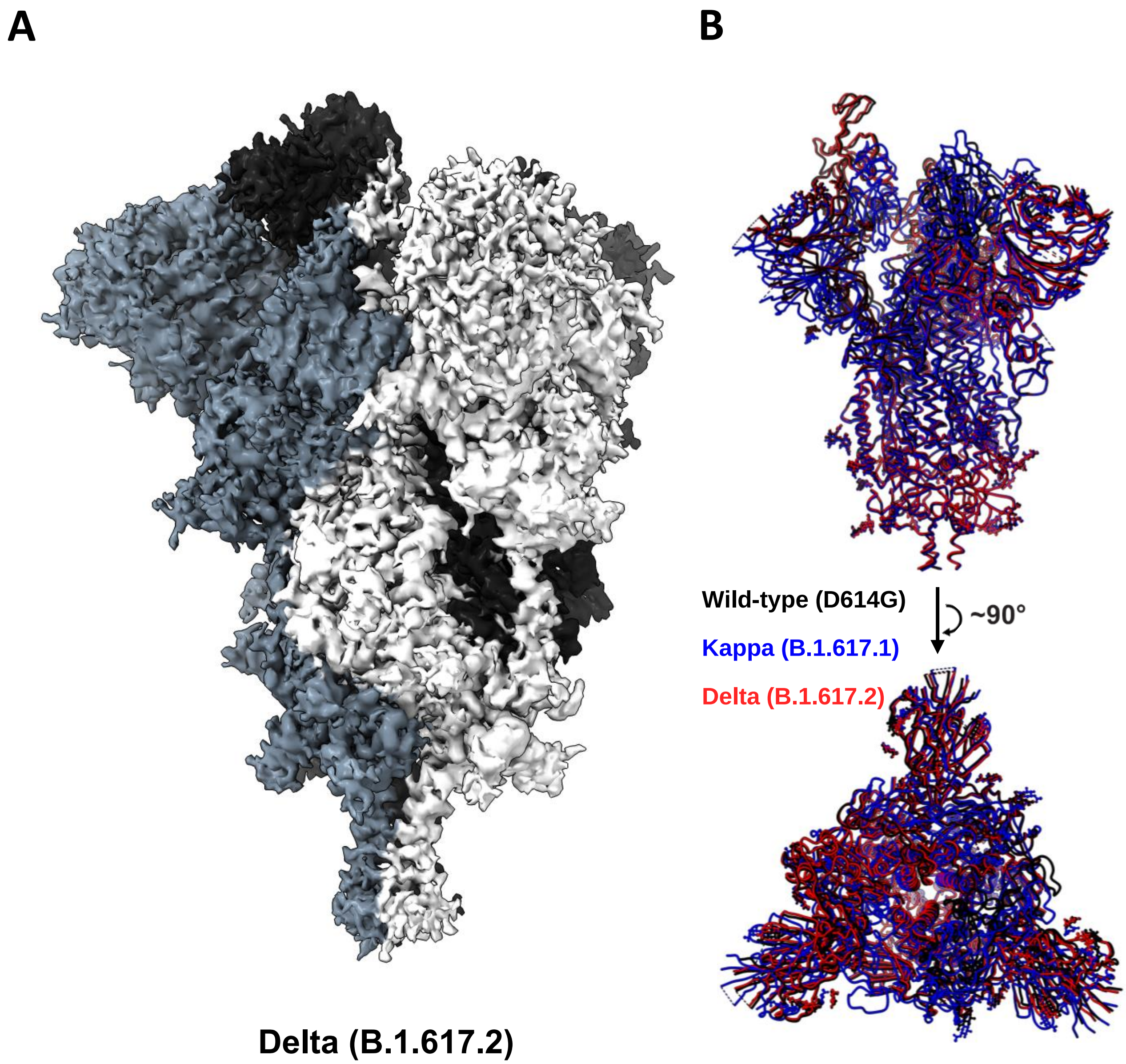


**Figure S11. CryoEm structure of the Delta (B.1.617.2) S protein ectodomain and its superposition with wild-type (D614G) and Kappa (B.1.617.1) structures. (A)** CryoEM density map of the Delta (B.1.617.2) variant S protein with each protomer coloured in greyscale. **(B)** Superposition of the wild-type (D614G – Black), Kappa (B.1.617.1 – Blue), and Delta (B.1.617.2 – Red) atomic models.

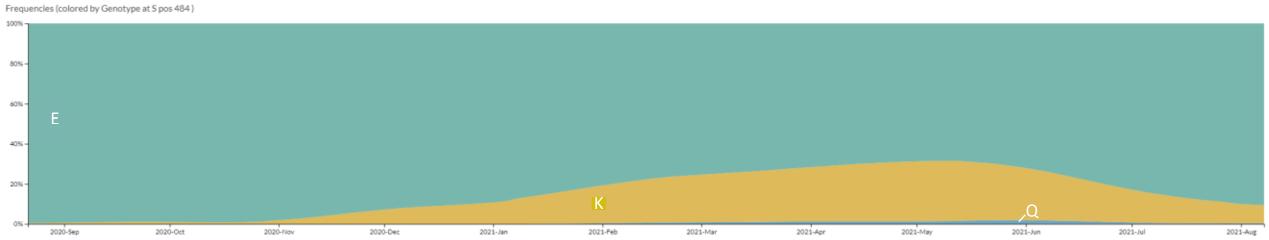


**Figure S12. Amino acid frequency at position 484 in global sequence deposits for the SARS-CoV-2 spike protein.** Residue frequency was derived from the Global Initiative on Sharing Avian Influenza Data (GISAID) database between September 2020 and August 2021.

**A**

| Sample | Sample Status | Vaccine Dose | Immunization to Serum Draw Time |
|---|---|---|---|
| P0 | Vaccine post-COVID19 | 1st | 4 Weeks |
| P1 | Vaccine | 1st | 3 Weeks |
| P3 | Vaccine | 1st | 6 Weeks |
| P5 | Vaccine | 1st | 1 Week |
| P6 | Vaccine | 1st | 3 Weeks |
| P8 | Vaccine post-COVID19 | 1st | 9 Weeks |
| P9 | Vaccine post-COVID19 | 1st | 7.5 Weeks |
| P10 | Vaccine post-COVID19 | 1st | 6.5 Weeks |
| P11 | Vaccine pre-COVID19 | 1st | 7 Weeks |
| P12 | Vaccine post-COVID19 | 1st | 8.5 Weeks |
| P13 | COVID19 | n/a | n/a (2 Weeks post-infection) |
| P14 | Vaccine | 2nd | 4 Weeks |

**B**

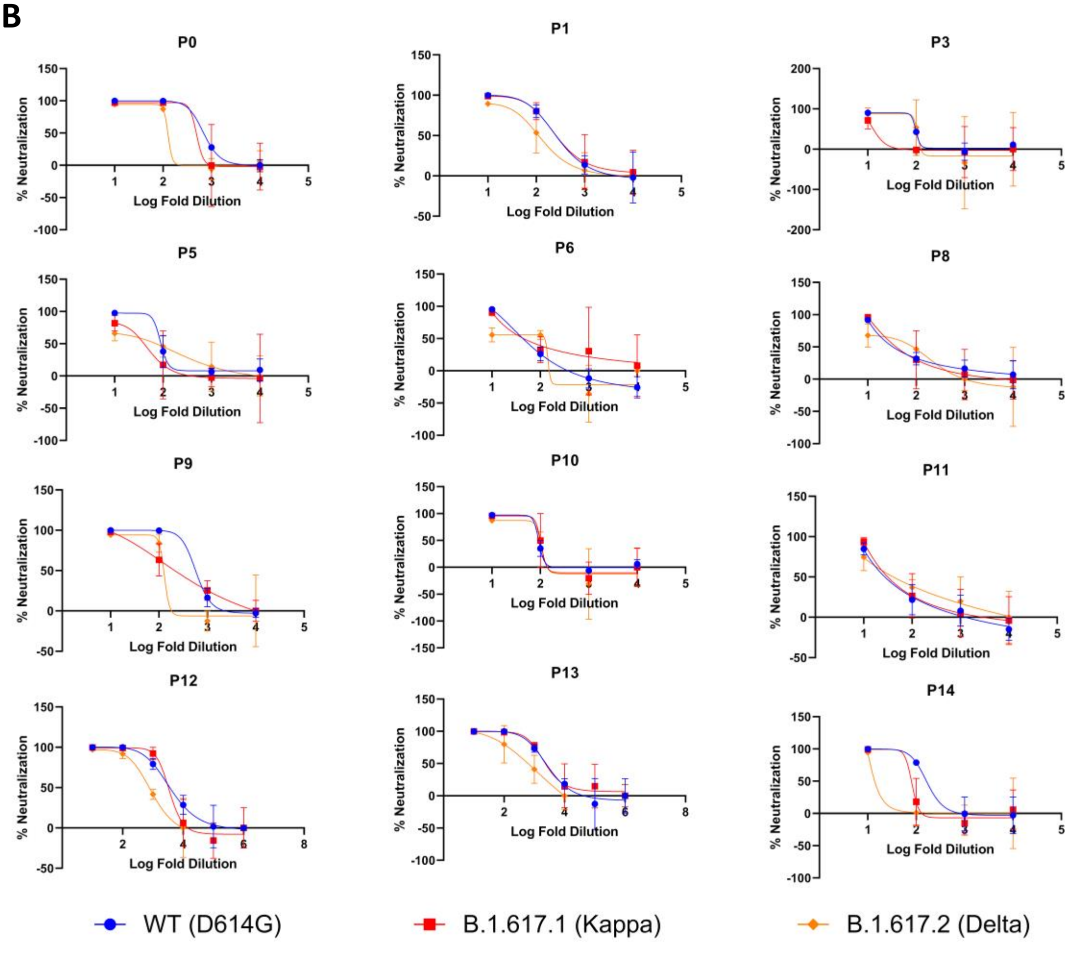


**Figure S13. Patient-derived sera sample information and raw pseudovirus neutralization data. (A)** Patient-derived sera sample information including sample number, vaccination status, vaccination dose number and immunization to serum draw time. **(B)** Raw data for the neutralization of Wild-type (D614G), Kappa (B.1.617.1), and Delta (B.1.617.2) S protein pseudotyped viruses with patient-derived sera samples.

### Table S1: CryoEM data collection, processing, refinement, and validation parameters for the structures reported in this publication.

| Structure | B.1.617.1 global refinement (EMDB 00000) (PDB XXXX) | B.1.617.1 focus refinement (EMDB 00000) (PDB XXXX) | B.1.617.1(Q484A) (EMDB 00000) (PDB XXXX) | B.1.617.1(Q484I) (EMDB 00000) (PDB XXXX) | B.1.617.1 + ACE2 global refinement (EMDB 00000) (PDB XXXX) | B.1.617.1 + ACE2 focus refinement (EMDB 00000) (PDB XXXX) | B.1.617.2 (EMDB 00000) (PDB XXXX) | B.1.617.2 + ACE2 global refinement (EMDB 00000) (PDB XXXX) | B.1.617.2 + ACE2 focus refinement (EMDB 00000) (PDB XXXX) |
|---|---|---|---|---|---|---|---|---|---|
| **Data collection** | | | | | | | | | |
| Microscope | Titan Krios G4 | | Titan Krios G4 | Titan Krios G4 | Titan Krios G4 | | Titan Krios G4 | Titan Krios G4 | |
| Detector | Falcon4 | | Falcon4 | Falcon4 | Falcon4 | | Falcon4 | Falcon4 | |
| Voltage (kV) | 300 | | 300 | 300 | 300 | | 300 | 300 | |
| Nominal magnification | 155,000 | | 155,000 | 155,000 | 155,000 | | 155,000 | 155,000 | |
| Defocus range (μm) | -3.0 to -0.5 | | -3.0 to -0.5 | -3.0 to -0.5 | -3.0 to -0.5 | | -3.0 to -0.5 | -3.0 to -0.5 | |
| Physical pixel (Å) | 0.5 | | 0.5 | 0.5 | 0.5 | | 0.5 | 0.5 | |
| Electron dose ($e^-/Å^2$) | 40 | | 40 | 40 | 40 | | 40 | 40 | |
| Exposure rate ($e^-/Å^2$/sec) | 24 | | 24 | 24 | 24 | | 24 | 24 | |
| Format of movies | EER | | EER | EER | EER | | EER | EER | |
| Number of raw frames | 399 | | 399 | 399 | 399 | | 399 | 399 | |
| Number of movies | 8,907 | | 6,750 | 8,380 | 8,478 | | 4,860 | 9,630 | |
| **Data processing** | | | | | | | | | |
| Number of fractions | 40 | | 40 | 40 | 40 | | 40 | 40 | |
| Number of extracted particles | 666,870 | | 592,644 | 585,964 | 619,611 | | 418,532 | 804,096 | |
| Number of particles for final map | 116,459 | | 261,710 | 76,565 | 105,037 | | 170,570 | 97,827 | |
| Symmetry imposed | C1 | C1 | C1 | C1 | C1 | C1 | C1 | C1 | C1 |
| Resolution (Å) | 3.16 | 3.01 | 2.25 | 3.62 | 3.02 | 3.27 | 2.25 | 3.27 | 3.52 |
| FSC threshold | 0.143 | 0.143 | 0.143 | 0.143 | 0.143 | 0.143 | 0.143 | 0.143 | 0.143 |
| **Refinement** | | | | | | | | | |
| Initial model used | 7MJG | 7MJG | 7MJG | 7MJG | 7MJM | 7MJN | 7MJG | 7MJM | 7MJN |
| Map sharpening B-factor ($Å^2$) | 53.3 | 53.7 | 43.4 | 37.6 | 50.1 | 76.6 | 35.1 | 42.9 | 94.0 |
| Composition (#) | | | | | | | | | |
| Atoms | 50,394 | 12,792 | 21,875 | 50,394 | 26,877 | 6,553 | 21,879 | 26,879 | 6,555 |
| Residues | 6,240 | 1,608 | 2,703 | 6,240 | 3,303 | 797 | 2,703 | 3,303 | 797 |
| Ligands | NAG:120 | NAG:12 | NAG:56 | NAG:120 | NAG:64 | NAG:7 | NAG:56 | NAG:64 | NAG:7 |
| B-factor ($Å^2$) | | | | | | | | | |
| Protein (min/max/mean) | 47.82/268.38/134.40 | 44.03/149.98/80.04 | 40.16/219.22/98.06 | 57.15/307.90/162.72 | 56.85/331.43/151.55 | 76.49/206.75/115.23 | 36.44/230.73/98.40 | 61.59/586.12/189.64 | 101.09/244.57/156.07 |
| Ligand (min/max/mean) | 102.29/288.20/169.64 | 76.17/133.82/96.25 | 62.03/224.10/111.43 | 120.49/336.15/207.56 | 88.95/288.56/152.96 | 116.95/149.58/132.83 | 56.80/234.34/112.22 | 94.99/565.69/175.62 | 164.13/210.83/180.46 |
| Bonds (RMSD) | | | | | | | | | |
| Length (Å) (# > 4σ) | 0.004 (0) | 0.005 (0) | 0.006 (0) | 0.005 (0) | 0.004 (0) | 0.005 (0) | 0.005 (0) | 0.004 (0) | 0.005 (0) |
| Angles (°) (# > 4σ) | 0.729 (14) | 0.835 (4) | 0.989 (31) | 1.017 (51) | 0.779 (8) | 0.885 (1) | 0.986 (24) | 0.769 (7) | 0.840 (1) |
| CC mask | 0.83 | 0.82 | 0.83 | 0.82 | 0.82 | 0.84 | 0.83 | 0.83 | 0.82 |
| **Validation** | | | | | | | | | |
| Ramachandran plot | | | | | | | | | |
| Residues favored (%) | 97.95 | 97.18 | 98.00 | 97.87 | 98.52 | 98.49 | 98.00 | 98.55 | 98.61 |
| Residues disallowed (%) | 0.00 | 0.00 | 0.00 | 0.00 | 0.03 | 0.00 | 0.00 | 0.00 | 0.00 |
| Rotamer outliers (%) | 0.33 | 0.00 | 1.10 | 0.04 | 0.07 | 0.71 | 0.97 | 0.21 | 0.14 |
| Clash score | 3.10 | 3.59 | 3.56 | 4.36 | 3.04 | 2.42 | 3.42 | 2.89 | 2.81 |
| MolProbity score | 1.11 | 1.30 | 1.18 | 1.25 | 1.09 | 1.02 | 1.13 | 1.08 | 1.07 |
